# Altered circadian rhythms and sleep in a new Angelman Syndrome mouse model

**DOI:** 10.1101/2021.10.26.465956

**Authors:** Shu-qun Shi, Carrie E. Mahoney, Pavel Houdek, Wenling Zhao, Matthew P. Anderson, Xinming Zhuo, Arthur Beaudet, Alena Sumova, Thomas E. Scammell, Carl Hirschie Johnson

## Abstract

Normal neurodevelopment requires precise expression of the key ubiquitin ligase gene *Ube3a*. Comparing newly generated mouse models for *Ube3a* down-regulation (models of Angelman syndrome) vs. *Ube3a* up-regulation (models for autism), we find reciprocal effects of *Ube3a* gene dosage on phenotypes associated with circadian rhythmicity, including the amount of locomotor activity. In contrast to previous reports, we find that *Ube3a* is imprinted in neurons of the suprachiasmatic nuclei, the pacemaking circadian brain locus. In addition, *Ube3a*-deficient mice lack the typical drop in wake late in the dark period and have blunted responses to sleep deprivation. Suppression of physical activity by light in *Ube3a*-deficient mice is not due to anxiety as measured by behavioral tests and stress hormones; quantification of stress hormones may serve as an easily measurable biomarker for evaluating potential therapeutic treatments for Angelman syndrome. We conclude that reduced *Ube3a* gene dosage affects not only neurodevelopment but also sleep patterns and circadian rhythms.

## Introduction

Although single-gene neurological disorders are uncommon, they are extremely valuable for investigating genetic and pathological mechanisms because their genetic underpinnings are usually more straightforward than in more complex genetic disorders [Shi et al., 2019]. Among monogenic neurodevelopmental disorders, Angelman syndrome (AS) is particularly interesting; deletion of only one copy (the maternal allele) of the ubiquitin ligase gene *Ube3a* confers haploinsufficiency because the paternal allele of *Ube3a* is imprinted and silenced in neurons of both AS and non-affected subjects [Jiang et al., 2010]. This haploinsufficiency leads to AS, resulting in mental disability, with impaired speech, delayed development, sleep disorders, epilepsy, and motor difficulties [Laan et al., 1999; Robb et al., 1989; Williams et al., 2006]. Interestingly, the paternal imprinting of *Ube3a* occurs only in neurons [Albrecht et al., 1997; Jiang et al., 1998; Weeber et al., 2003], and not in glia or non-neural peripheral tissues [Dindot et al., 2008; Gustin et al., 2010; Judson et al., 2014; Nakao et al., 1994]. In healthy subjects, the maternal copy of *Ube3a* is active in adult neurons while the paternal copy is silenced. On the other hand, extra copies of *Ube3a*, as in duplications and triplications of the region of chromosome 15 that harbors *Ube3a* are a common large genetic alteration that is linked to autism spectrum disorder (ASD) [Sutcliffe et al., 1997; Lehman et al., 2009]. Therefore, appropriate gene dosage of *Ube3a* is critical for normal neural development–the equivalent of one active allele allows normal development, whereas too little leads to AS, and too much leads to autism.

About 50-75% of individuals with AS have a sleep disorder, such as short sleep duration and increased sleep onset latency, and these sleep disruptions are one of the syndrome’s most stressful problems for families caring for a person with AS [Goldman et al., 2012; Pelc et al., 2008; Smith et al., 1996]. These sleep phenotypes in AS could be due to a misalignment of the circadian system, as the timing of sleep is regulated by the circadian clock [Scammell et al., 2017; Schwartz et al., 2019]. While most research on sleep disruptions in AS is limited to clinical/behavioral observations, one study of daily profiles of the hormone melatonin concluded there was a high prevalence of circadian rhythm sleep disorders in AS [Takaesu et al., 2012].

Though clinical research on sleep in AS is limited, animal models provide many opportunities to identify the impact of *Ube3a* deficiency on the circadian and sleep systems. The two currently available mouse models for AS include an exon 5-specific deletion within *Ube3a* that also creates a frame shift (herein called Ud5-m-/p+ for deletion of exon 5 from the transcriptional start site in the maternal *Ube3a* allele combined with an intact paternal allele) [Jiang et al., 1998]. The second model is caused by a larger (1.6 Mb) chromosomal deletion that removes the maternal copies of *Ube3a* as well as the adjacent *Atp10a* and *Gabrb3* genes (herein called UG-m-/p+)[Jiang et al., 2010]. These mouse models demonstrate phenotypes similar to those seen in human AS patients, including motor dysfunction, impaired locomotion, seizures, learning deficits, abnormal EEG (electroencephalogram) patterns, and changes in CaMKII phosphorylation that reduce hippocampal long-term potentiation (LTP)[Colas et al., 2005; Jiang et al., 1998; Shi et al., 2015; Weeber et al., 2003; Woerden GM et al., 2007]. Most people with AS (∼75%) have large deletions of the maternal chr15q11-13 alleles, including *Ube3a, Atp10a*, and *Gabrb3*, and therefore the larger-deletion UG-m-/p+ mouse model is generally considered to better reflect AS in humans than the smaller U-m-/p+ deletion model [Jiang et al., 2010]. On the other hand, the Ud5-m-/p+ model is specific for the *Ube3a* gene that is primarily responsible for AS and therefore allows precise association of aberrant phenotypes with reduced dosage of *Ube3a* [Jiang et al., 1998] (but see the paragraph below concerning inconsistencies with the Ud5-m-/p+ model).

We have reported that the robustness of circadian rhythms is diminished in both Ud5-m-/p+ and UG-m-/p+ mice as assessed by longer free-running periods in constant darkness (DD), accelerated phase resetting, and greater suppression of activity in constant light (LL)[Shi et al., 2015]. Another research group has reported that Ud5-m-/p+ mice have a reduced capacity to accumulate sleep pressure, both during their active period and in response to forced sleep deprivation [Ehlen et al., 2015]. Contrary to predictions, they also reported that the neurons of the suprachiasmatic nuclei (SCN, the site of the master circadian clockwork in mammals) were unusual in that they expressed more UBE3A protein than expected, and therefore the paternal allele of *Ube3a* was exceptionally <u>not</u> imprinted in the central clock neurons of Ud5-m-/p+ mice [Ehlen et al., 2015; Jones et al., 2016]. This observation was particularly surprising because the circadian period is lengthened in cultured SCN brain slices of Ud5-m-/p+ mice isolated *in vitro* [Shi et al., 2015] and that result is difficult to explain if UBE3A levels were unchanged.

Those disparate observations were made with the Ud5-m-/p+ mouse model [Ehlen et al., 2015; Jones et al., 2016; Shi et al., 2015], which has been the standard mouse model of AS. However, multiple labs have reported challenges with the Ud5-m-/p+ model such as strain dependency and loss of phenotype over time [Born et al., 2017; Huang et al., 2013; Dodge’2020]. Moreover, the Ud5-m-/p+ mouse model may not be a complete null mutation of *Ube3a* because the deleted exon 5 is subject to brain-specific alternative splicing that can result in a truncated *Ube3a* transcript (Dr. Scott Dindot, personal communication, September 2021). This truncated mRNA is not translated and its functions, if any, are unknown. These inconsistencies have led to the generation of new animal models for AS which could better reflect the human AS phenotype, including a model in rats [Dodge et al., 2020]. To take full advantage of the genetics that mouse models enable, it is essential to develop a mouse line that specifically targets the *Ube3a* gene with a consistent phenotype similar to Angelman syndrome. Because a larger deletion within the *Ube3a* gene should more closely mimic the condition in people with AS, we studied the circadian and sleep phenotypes of a new mouse model, i.e., Ud6-m-/p+ mice that harbor a larger deletion of *Ube3a’*s exon 6 (a 1247 nt deletion that also confers a frame shift, whereas the deletion in exon 5 of the Ud5-m-/p+ mouse is only 290 nt, Fig. 1A). We found that phenotypes of this new model recapitulate, confirm, and extend the observations previously made with the Ud5-m-/p+ model for circadian [Shi et al., 2015] and sleep behavior [Ehlen et al., 2015]. Moreover, and in contrast to previous studies [Ehlen et al., 2015; Jones et al., 2016], we find that the expression of UBE3A in the SCN to a degree that is consistent with paternal imprinting, and therefore the neurons within the SCN are not an exception to the conclusion that the paternal allele of *Ube3a* is imprinted within neurons.

**Figure 1.**
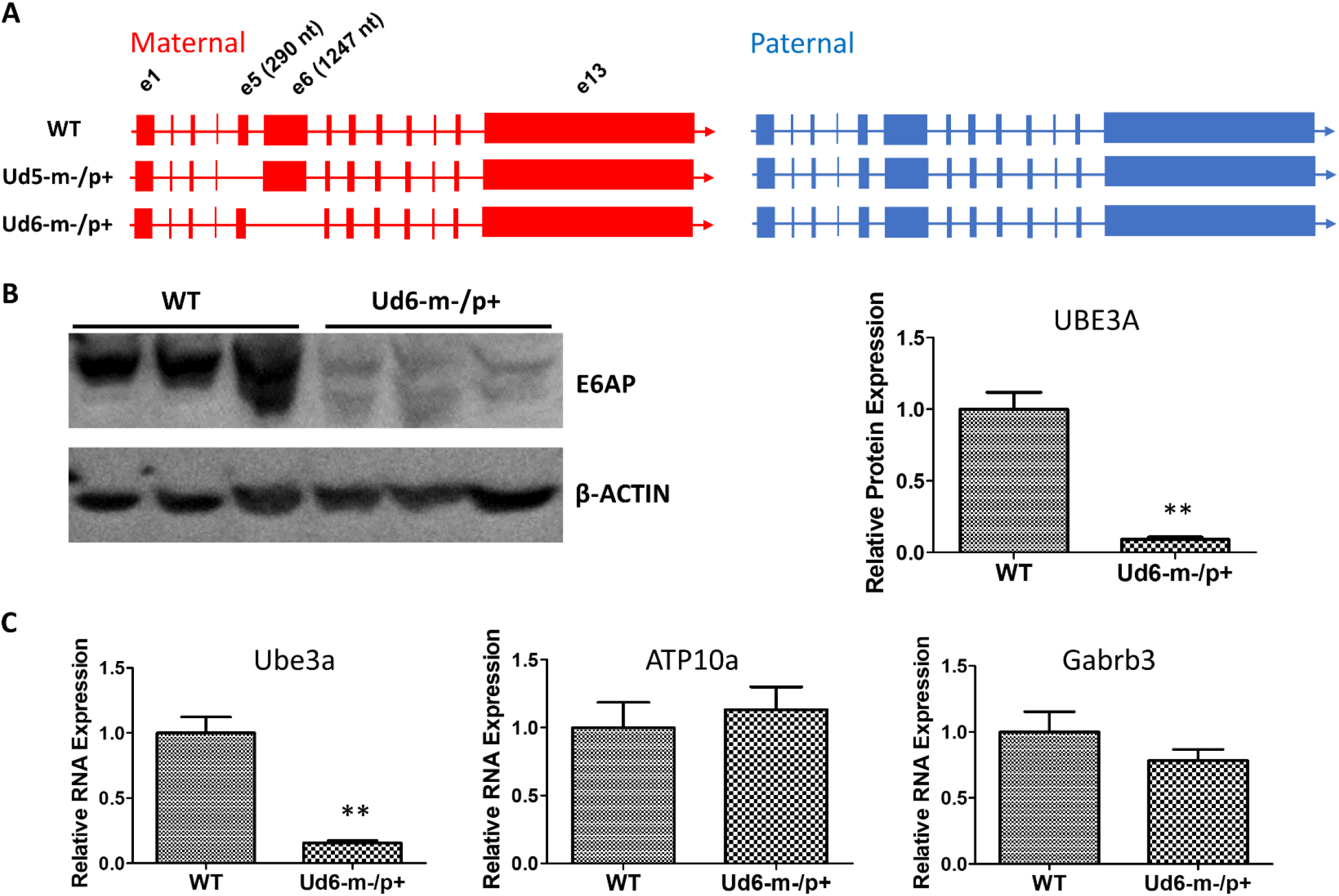
Ube3a is imprinted in the brain of the Ud6-m-/p+ mouse. (A) Female mice harboring one *Ube3a* knockout allele (left panel) were crossed with wild-type background (WT C57BL/6J) male mice (right panel) to obtain the maternal deletion Angelman model mice. The maternal and paternal *Ube3a* allele arrangements are WT (upper), the original Angelman model mouse strain harboring an exon 5 deletion in Ube3a gene, Ud5-m-/p+ (middle), and the new strain knockout of exon 6 in the Ube3a gene, Ud6-m-/p+ (lower). (B) **Left panel**, immunoblots for UBE3A protein in the cerebellum of Ud6-m-/p+ and WT mice. β-Actin serves as the loading control (each lane comes from a separate mouse; one representative immunoblot is shown). **Right panel**, densitometric analyses of the immunoblot (The expression of WT is normalized to 1.0, n=3 for both WT and Ud6-m-/p+). ** p < 0.01 by two-tail unpaired T test. (C) *Ube3a, ATP10a*, and *Gabrb3* mRNA transcript abundances in cerebellum of Ud6-m-/p+ and WT mice were quantified by real-time PCR (n=3 for both WT and Ud6-m-/p+). ** p < 0.01 by two-tail unpaired T test.

## Results

### Reduction of UBE3A expression in the Ud6-m-/p+ brain

To confirm that the levels of UBE3A protein were down-regulated by imprinting of the paternal allele in the Ud6-m-/p+ mouse, we measured UBE3A levels in the cerebellum of brains by immunoblotting with an anti-UBE3A antibody. In Ud6-m-/p+ mice, UBE3A protein was reduced to only 10% of the WT level (Fig. 1B). Consistent with the lower gene expression indicated by minimal UBE3A protein levels, the *Ube3a* transcript abundance in the cerebellum of Ud6-m-/p+ mice was only 15% that of the WT levels. At the same time, mRNA levels for the *Atp10a* and *Gabrb3* genes did not differ between WT and Ud6-m-/p+ mice (Fig. 1C). The *Atp10a* and *Gabrb3* genes are immediately adjacent to *Ube3a* on mouse chromosome 7 and are not imprinted; therefore, they serve as controls for the specificity of down-regulation of *Ube3a* in the Ud6-m-/p+ mouse.

To explore and visualize the spatial expression of Ube3a expression in two key areas of the brain, we conducted immunohistochemical (IHC) localization of UBE3A in the hippocampus and suprachiasmatic nuclei (SCN, the site of the central mammalian biological clock mechanism [Reppert & Weaver., 2002]). We detected UBE3A immunopositivity by standard avidin-biotin methods, or by fluorescent dyes for antigen co-localization [Liska et al., 2021; Sumova et al., 2002]. The IHC results were quantified by measuring optical density as a measure of the expressed protein levels in the hippocampal CA1, CA3, and DG regions, where the cells are in tight proximity (cell bodies are dense and difficult to reliably resolve), and by calculating the number of individual immunopositive cells in the SCN (cell number was used for the SCN because of its unique anatomy and structure that allows cells bodies to be resolved). The optical density of UBE3A staining was abundant in all three hippocampal regions of the WT mice, but it was below the detection limit in Ud6-m-/p+ mice (Fig 2A). UBE3A expression was consistently reduced across the entire hippocampus including the CA1, CA3, and DG regions (ANOVA, p < 0.0001 for all regions, Fig 2B) that are important regions for functional learning and memory [Dodge et al., 2020; Jiang et al., 1998]. Therefore, Ube3A protein expression was silenced in the hippocampus of Ud6-m-/p+ mice.

**Figure 2.**
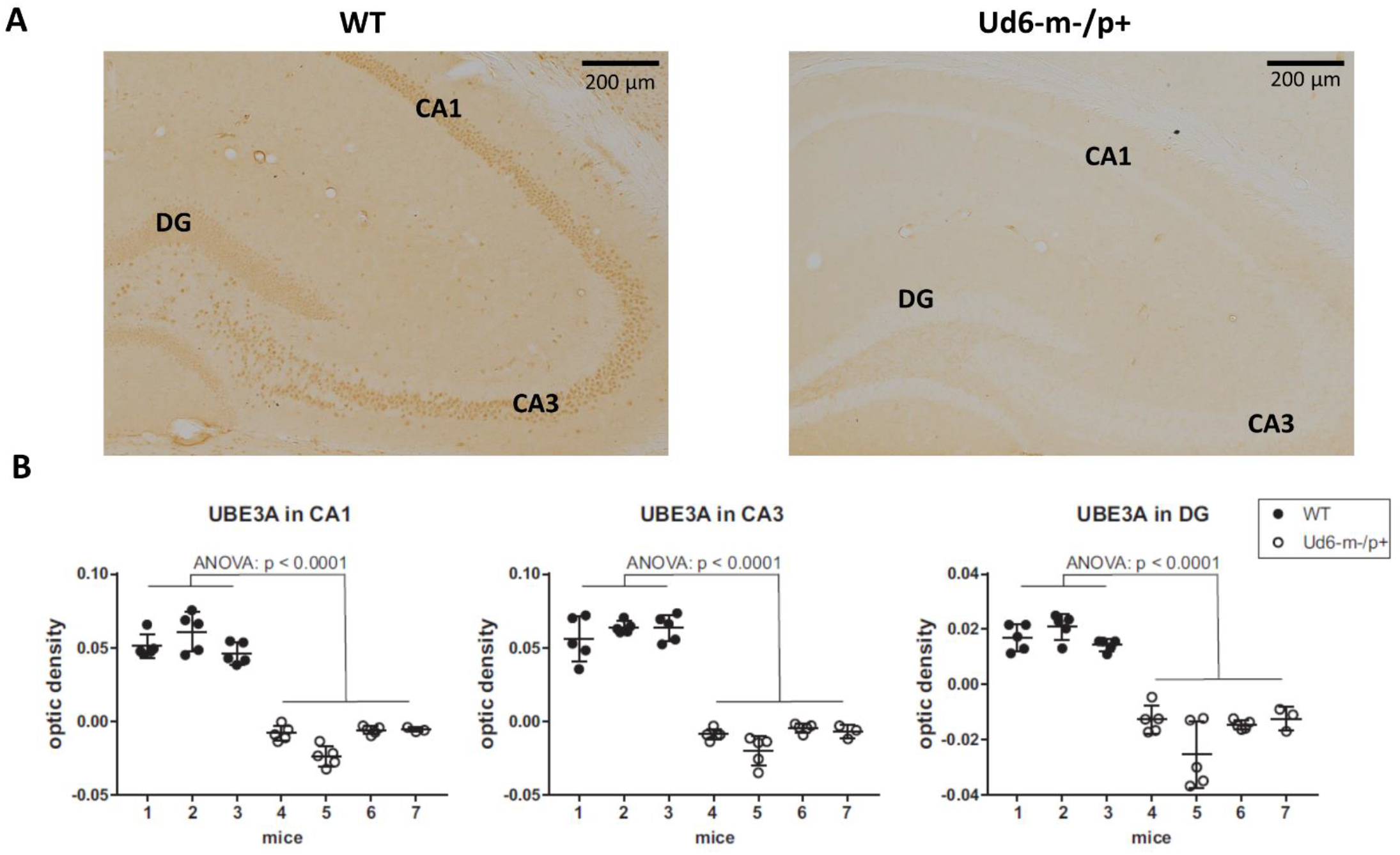
Reduced abundance of UBE3A protein in the Ud6-m-/p+ hippocampus. (**A**) Representative images of immunohistochemical (IHC) detection of UBE3A in coronal brain sections through the hippocampal CA1-3 and dentate gyrus (DG) of wild type (WT; left) and Ud6-m-/p+ (Ud6; right) mice. UBE3A-immunopositive cells (brown dots) are absent in all hippocampal regions of Ud6-m-/p+ mice. Scale bar = 200 µm. (**B**) Quantification of IHC results by optical density measurement in the subregions of the hippocampus. Data are expressed as individual values for each separate section (n = 3-5 per brain) and as the group mean ± S.D., WT n=3, Ud6-m-/p+ n=4 (individual mice numbered along abscissa; numbers 1 - 3 for WT and 4 – 7 for Ud6-m-/p+).

Additionally, we found robust suppression of UBE3A-immunopositivity in the SCN of Ud6-m-/p+ mice (Fig. 3A). Ud6-m-/p+ mice had only about 25% of the number of SCN cells expressing UBE3A (Fig. 3B, ANOVA; p < 0.0001) compared to the SCN in WT mice (Fig. 3C, t-test; p < 0.0001). By examination of confocal images under high magnification, we confirmed that *Ube3a* is expressed in neurons but not in astroglia within the SCN and hippocampus because UBE3A co-localized with ELAVL4, a marker for neurons [Jung and Lee., 2021], but not with GFAP, a marker for astrocytes [Brenner et al., 1994] (Supplemental Fig. S1). In contrast to a prior study of the Ud5-m-/p+ mouse [Jones et al., 2016], we clearly confirm that UBE3A is down-regulated in the SCN of Ud6-m-/p+ mice to an extent that is entirely consistent with imprinting of the paternal *Ube3a* allele in SCN neurons. If the paternal allele of *Ube3a* were fully active, there should be 50% of UBE3A expression as compared with WT because of the deletion of the maternal allele; however, with our criterion that is based upon the number of UBE3A cells, we find that 25.5% of SCN cells retain some UBE3A expression (Fig. 3C). Detailed inspection of the overlap between UBE3A and ELAVL4 or GFAP staining revealed that the majority of the UBE3A-positive cells were neurons but there were a few cells in the SCN sections that expressed UBE3A but not ELAVL4 or GFAP (not shown).Therefore, some of these UBE3A-positive cells within the SCN of Ud6-m-/p+ mice may be non-neuronal.

**Figure 3.**
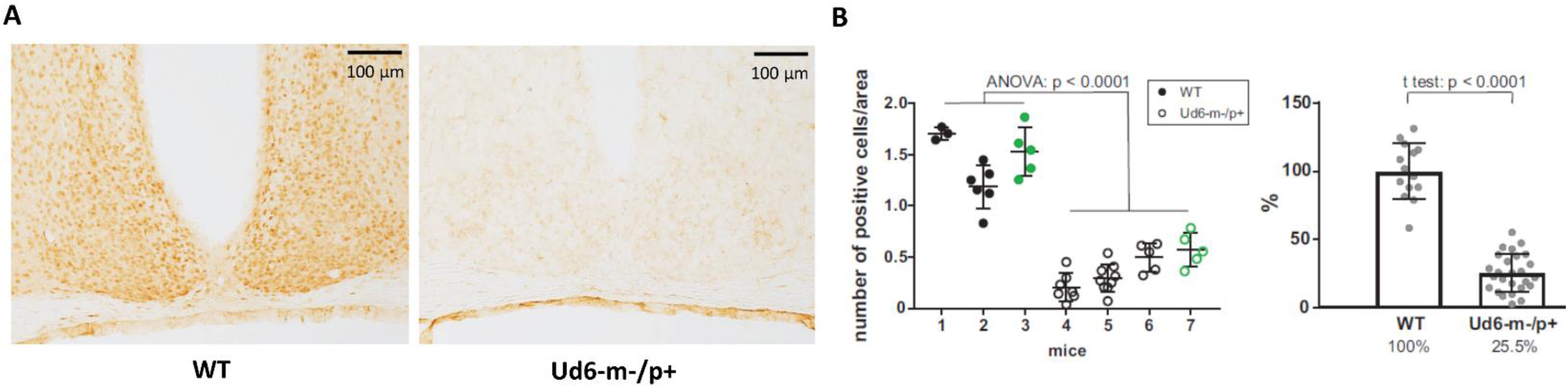
Reduced abundance of UBE3A-positive neurons in Ud6-m-/p+ SCN. (**A**) Representative IHC images of SCN sections from wild type (WT; left) and Ud6-m-/p+ (Ud6; right) brains. The UBE3A-immunopositive cells (brown dots) are densely distributed throughout the SCN in WT, but are sparse in Ud6. Scale bar = 100 µm. (**B**) Quantification of positively stained cells in the SCN. Data are expressed as the number of positive cells calculated in 3-9 separate sections per each brain, and the group mean ± S.D. (individual mice numbered along the abscissa; numbers 1 - 3 for WT and 4 – 7 for Ud6-m-/p+). Black dots represent immunohistochemical diaminobenzidine staining (WT n=2, Ud6-m-/p+ n=3), while green dots represent data from immunofluorescence staining (WT n=1, Ud6-m-/p+ n=1; see Supplemental Figure 1). (**C**) Combination of data from panel **B** depicting the relative UBE3A staining in all SCN sections where 100% is set as the mean of all WT values (WT n=3, Ud6-m-/p+ n=4).

### Altered rotarod and marble-burying phenotypes in Ud6-m-/p+ mice

The previously generated mouse models of Angelman syndrome (Ud5-m-/p+ and UG-m-/p+) exhibit altered phenotypes in rotarod and marble-burying behavioral tests [Jiang et al.,1998; Jiang et al., 2010; Sonzogni et al., 2018]. We used the accelerating rotarod to study motor coordination and motor learning skills and confirmed that the Ud6-m-/p+ mice also performed significantly worse on the rotarod than their WT littermates (p < 0.0001 for the genotype factor by two-way ANOVA). There was a clear sex difference in the improvement in fall-off time over the trial testing, with females exhibiting better motor learning over trials in the task than males (Fig. 4A). AS model mice also have altered marble burying behavior, which is a test used to assess repetitive and perseverative behavior as well as anxiety [Sonzogni et al., 2018]. We found that Ud6-m-/p+ mice bury fewer marbles than WT mice when tested both in the day (ZT 3-5) and at night (ZT 15-17), indicating reduced anxiety and compulsiveness [Angoa-Perez et al., 2013; Thomas et al., 2009] (p < 0.0001 for the genotype factor by two-way ANOVA, Fig. 4B). We also confirmed that the repetitive and perseverative behavior as well as anxiety tested by marble burying has a significant day and night variation in WT animals (p=0.004 by T test, Fig. 4B), suggesting an importance of circadian factor in preclinical study. Therefore, consistent with other studies, the Ud6-m-/p+ model mice exhibit altered behaviors as compared with WT littermates.

**Figure 4.**
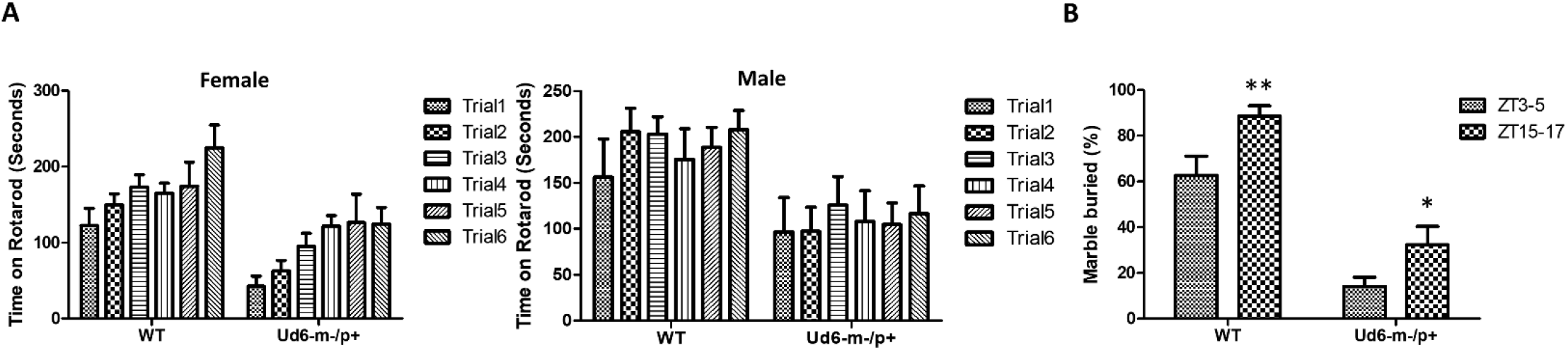
Rotarod & Marble burying behaviors in Ud6-m-/p+ mice. **(A)** Accelerating rotarod data for Ud6-m-/p+ vs. WT mice, tested over two contiguous days for three trials per day at ZT8-11 under LD conditions. The data are stratified by trials and the sex of mice (Male: 6 WT and 5 Ud6-m-/p+; Female: 8 WT and 11 Ud6-m-/p+). mean ± SEM. Female (8 WT and 11 Ud6-m-/p+): Genotype p < 0.0001 and time p < 0.0001 by two-way ANOVA. Male (6 WT and 5 Ud6-m-/p+): Genotype p < 0.05 and time p > 0.05 by two-way ANOVA. **(B)** Marble burying behavior tested both during the day (ZT3-5) and at night (ZT15-17) under LD conditions (WT: N=6, Ud6-m-/p+: N=10). mean ± SEM. p<0.0001 for both the genotype factor (WT vs. Ud6-m/p+) and the time factor (ZT3-5 vs. ZT15-17) by two-way ANOVA. ** p<0.01 for WT and * p<0.05 for Ud6-m-/p+ assayed at ZT3-5 and ZT15-17 by the Bonferroni posttest.

### Haploinsufficiency of *Ube3a* alters locomotor activity patterns in response to LL

Environmental exposure to constant light (LL) disrupts and/or suppresses circadian activity patterns in mice, and we previously reported an enhanced effect of LL on Ud5-m-/p+ mice [Shi et al., 2015]. We observed a similar LL-dependent effect on Ud6-m-/p+ mice as compared with WT mice (Fig. 5). During the exposure to LL, Ud6-m-/p+ mice displayed dramatically reduced activity levels and suppressed rhythms of wheel-running as compared with WT mice; most of Ud6-m-/p+ mice (5 out of 8) were completely arhythmic and displayed lower activity under LL (Fig. 5A). The supression of running-wheel activity largely recovered after mice were transferred from LL to DD (Fig. 5A, left & middle panels). In addition, Ud6-m-/p+ mice had less general locomotion in both LL and after transfer from LL to DD as assessed by infrared sensors (Fig. 5B left & middle panels). Moreover, the shortening of the circadian period of wheel-running behavior upon transfer from LL to DD was smaller in Ud6-m-/p+ than in WT mice (Fig. 5A, right panel). While exposed to LL, the circadian periods of WT and Ud6-m-/p+ mice were comparable for both running-wheel activity (Fig. 5A right) and for general activity (Fig. 5B right), but after transfer to DD, the period of WT mice shortened ∼1.5 h (1.48±0.33 h for running wheels, 1.45±0.37 h for general activity, right panels of Fig. 5), whereas the LL to DD transfer evoked only ∼1.0-1.1 h shortening in Ud6-m-/p+ mice (0.99±0.89 h for running wheels, 1.14±0.33 h for general activity, right panels of Fig. 5). The observation that LL-suppression of circadian amplitude is magnified while light-dependent period changes are diminished in Ud6-m-/p+ mice further supports our previous conclusion that normal levels of UBE3A expression enhance circadian oscillator strength, and therefore that Angelman mice have weaker circadian pacemakers [Shi et al., 2015]. In addition to altered circadian locomotor behavior in LL, the free running period of Angelman mice in DD trended towards longer periods when measured by rotating-saucer exercise wheels (Fig. S2). Together with our previously reported observations of Ud5-m-/p+ mice [Shi et al., 2015], the data of Fig. S2 also imply a sex difference in the magnitude of period lengthening in Angelman mouse models.

**Figure 5.**
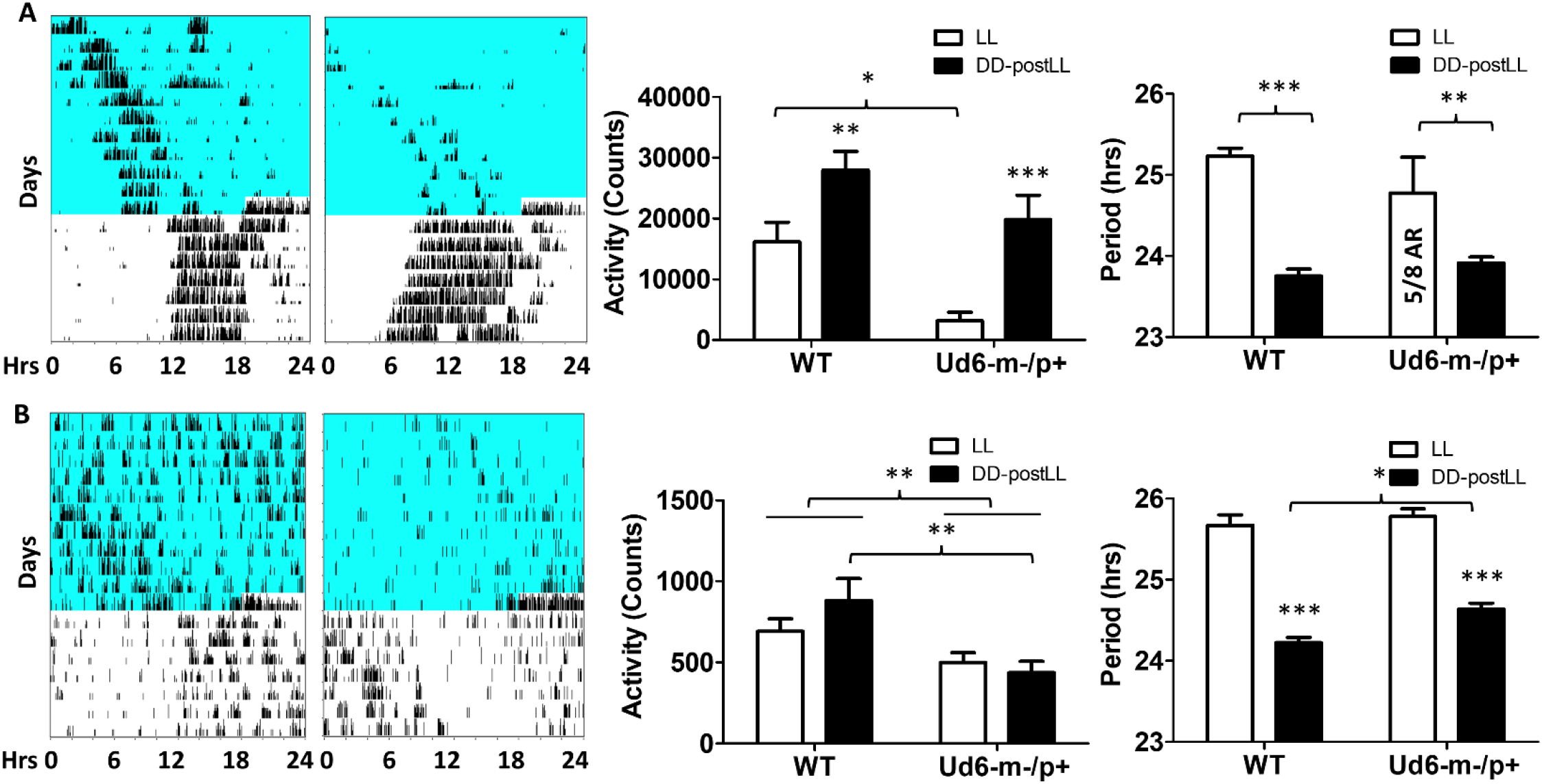
Light suppresses locomotor activity in Ud6-m-/p+ mice. (A) : **Left**: Representative wheel-running activity records (actograms) of mice are shown in the double-plotted format. Mice were initially in constant light (LL, blue background indicates light) and were then transferred to constant darkness (DD, white background). Total daily wheel-running activity (**middle panel**) and period analysis (**right panel**) during LL and after transfer from LL to DD (“DD-postLL”) were quantified (WT: n=8, Ud6-m-/p+: n=8). Data are plotted as mean ± SEM and analyzed by two-way ANOVA. Wheel-running activity (middle panel): p<0.05 of the genotype factor (WT vs. Ud6-m/p+) and p<0.001 of the timing factor (LL vs. DD-postLL), *p<0.05, ** p<0.01, and ** p<0.001 by the Bonferroni posttest as indicated. Wheel-running activity period (right panel): p<0.001 of the timing factor (LL vs. DD-postLL), ** p<0.01, and ** p<0.001 by the Bonferroni posttest as indicated. (B): **Left**: Representative total activity was monitored by infrared motion sensors in LL and DD from WT and Ud6-m-/p+ mice. Total daily activity by infrared sensors (**middle panel**) and period analysis (**right panel**) during LL and DD-post LL were quantified (WT: n=6, Ud6-m-/p+: n=6). Data are plotted as mean ± SEM and analyzed by two-way ANOVA. Infrared activity (middle panel): p<0.01 of the genotype factor (WT vs. Ud6-m/p+), p<0.01 by the Bonferroni posttest as indicated. Infrared activity period (right panel): p<0.001 of the timing factor (LL vs. DD-postLL), *** p<0.001 by the Bonferroni posttest for LL vs. DD-PostLL, * p<0.05 as indicated.

### Additional UBE3A has opposite effects to UBE3A deficiency on clock properties

Our observations of Ud6-m-/p+ mice in this study and of Ud5-m-/p+ mice in our previous study [Shi et al., 2015] indicate that the reduced expression of UBE3A in neurons leads to longer circadian periods and altered behavior in the light. What about enhanced expression of UBE3A? A simple prediction would be that additional expression of UBE3A should shorten the circadian period, and this is what we observed. Using ubiquitous overexpression of FLAG-*Ube3a* in a BAC transgenic mouse model for autism [Krishnan et al., 2017; Smith et al., 2011], we found that the circadian period of the SCN *in vitro* was shortened by the enhanced expression of *Ube3a* compared with that in WT SCN from the same mixed background (Fig. 6A). Ectopic UBE3A expression in the SCN from 1x Tg mice was assessed by immunoblots, confirming that one additional copy of the *Ube3a* gene introduced by the FLAG-*Ube3a* transgene increased levels of UBE3A protein (Fig. 6B). Moreover, while UBE3A deficiency in Angelman models suppresses the activity levels of mice in the light interval of LD (Fig. 4 and [Shi et al., 2015]), we found that *Ube3a* transgenic mice run more on wheels during the light/day phase of LD when the mice are usually inactive/asleep (Fig. 6C&D), and this increased activity is *Ube3a* gene dosage-dependent; 2xTg mice displayed twice as much activity in the light interval as did 1xTg mice (Fig. 6D).

**Figure 6.**
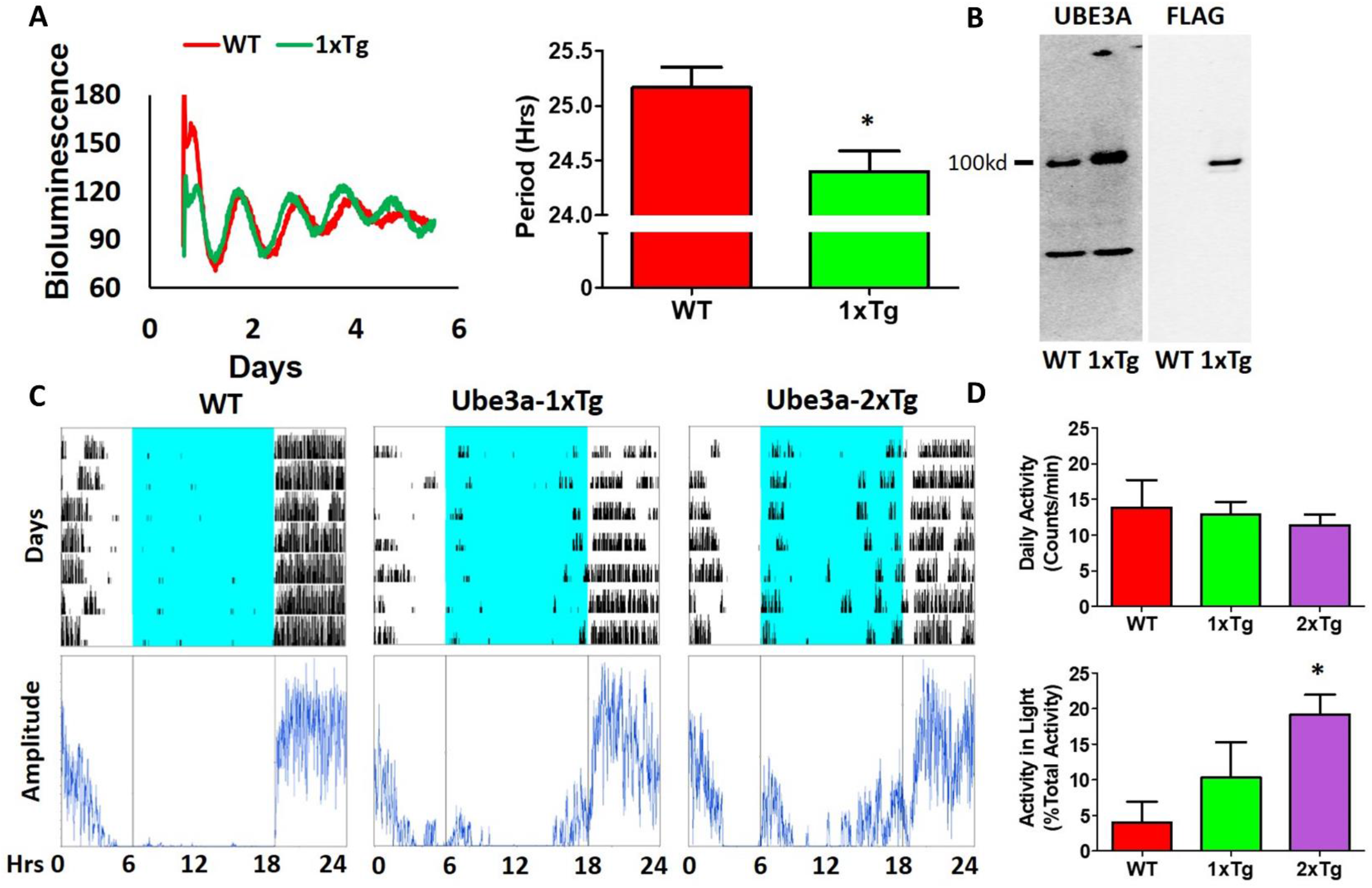
Additional expression of *Ube3a* shortens molecular rhythms in SCN slices *in vitro* and increases light-phase activity *in vivo*. (A) : Extra dosage of the *Ube3a* gene (as the *Ube3a*-FLAG BAC transgene, aka 1x Tg) alters molecular rhythms in explanted suprachiasmatic nuclei (SCN) slices, as assessed with a luminescence reporter of mPER2 expression. SCN slices were collected from P_mPer2_::mPER2-LUC knockin mice in the WT and 1x Tg backgrounds. Tissue explants were dissected on Day 0, recorded with a LumiCycle apparatus, and luminescence traces are shown in the left panel. Red: WT; Green: *Ube3a* 1x transgene. **Histograms**: period analysis of the SCN data plotted as mean ± SEM (WT: n= 3, 1x Tg: n=4), *p<0.05 by two-tail unpaired T test. (B) : Immunoblots of UBE3A and FLAG tag in the SCN of WT and 1x Tg mice. The arrow points to the specific bands of UBE3A-FLAG identified by anti-UBE3A and anti-FLAG antibodies, respectively. (C) : Representative data for mice in LD depict the wheel-running activity (upper panels) of WT (no additional *Ube3a* dosage), 1x Tg (one copy of transgenic *Ube3a*-FLAG), and 2x Tg (two copies of transgenic *Ube3a*-FLAG) with accompanying amplitude analyses (lower panel) computed by ClockLab. (D) : Wheel-running activity analyses for total daily activity (upper panel) and activity in the light phase (lower panel) were computed by ClockLab software for the data under the protocol of panel C. Data are plotted as mean ± SEM (WT: n=3, 1x Tg: n=4, 2x Tg: n=3), *p<0.05 between WT and 2x Tg by two-tail unpaired T test.

### Reduced stress hormones and anxiety in Ud6-m-/p+ mice

The reduced activity of Ud6-m-/p+ mice in LL could be interpreted as an enhanced stress response to illumination. To rule out the possibility that this altered behavior was caused by stress or anxiety, we measured stress hormones and neurotransmitters in the serum and brain during the day phase of LD. Corticosterone and norepinephrine release are regulated by stress and the circadian clock [Le Minh et al., 2001; Ikegami et al., 2020]. In Ud6-m-/p+ mice, we found that serum corticosterone levels are lower at ZT10-12 (Fig. 7A), a peak phase for this hormone in the blood. Content of another stress-related hormone, norepinephrine, is similarly reduced in the hypothalamus of Ud6-m-/p+ mice at ZT 10-12 (Fig. 7A). We also found a reduced norepinephrine level in the cerebellum of Ud5-m-/p+ mice at the same early night phase (Fig. S3). In contrast, deficiency of *Ube3a* expression did not alter the levels of other key neurotransmitters (GABA, DA, DOPAC, and 5-HT) in the hypothalamus or the cerebellum (Fig. 7B & Fig. S3). These decreased corticosterone and norepinephrine levels are consistent with the aforementioned results from the marble burying tests (Fig. 4B) towards indicating a reduced behavioral anxiety in the Ud6-m-/p+ mice. Therefore, the anxiety/stress levels of Ud6-m-/p+ mice are less than–not greater–than those of WT mice, which is similar to the conclusions of studies with Angelman human subjects [Robb et al., 1989], and consequently the reduced activity of Angelman model mice is probably not due to light-induced anxiety.

**Figure 7.**
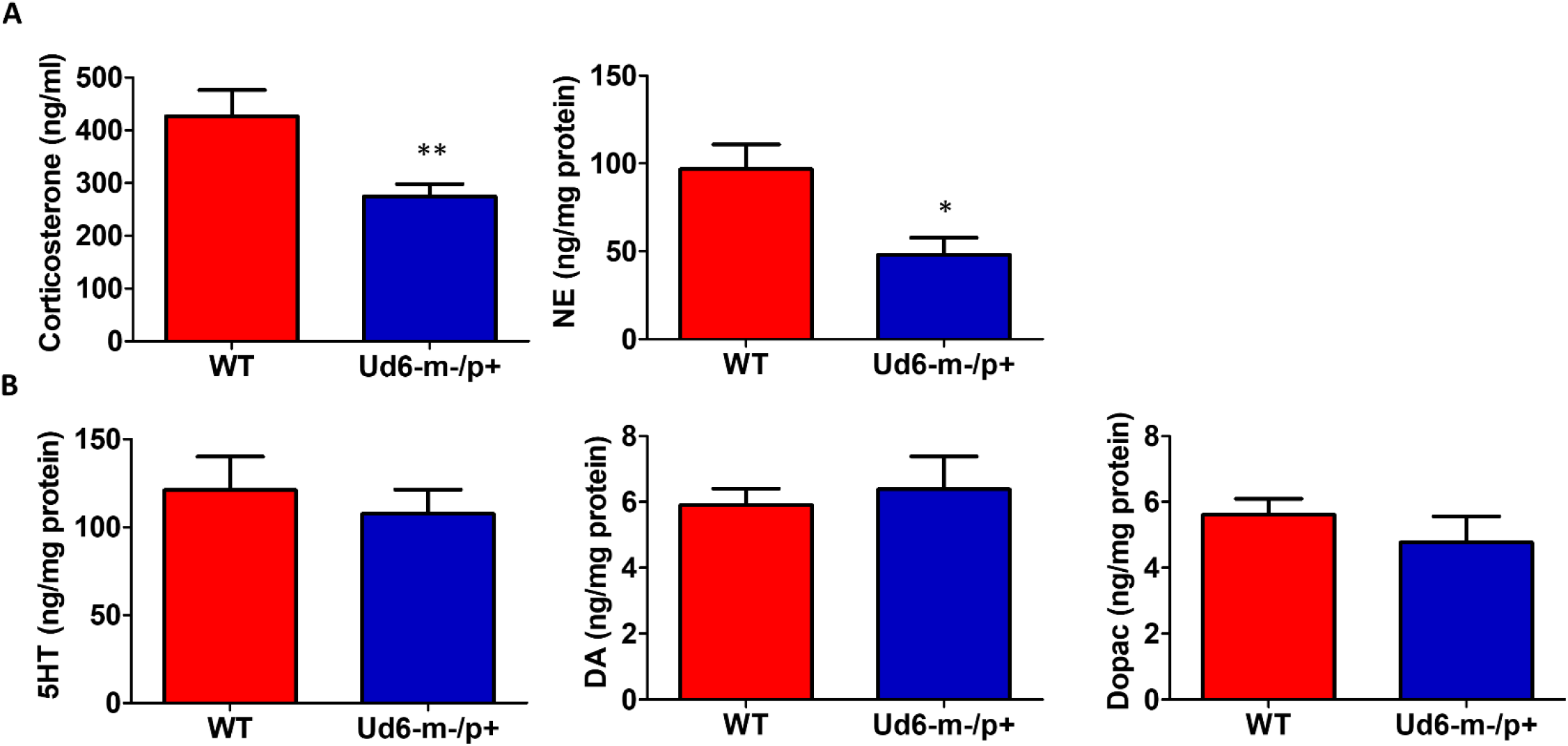
Reduced anxiety and stress hormones in Ud6-m-/p+ mice. (A) Serum corticosterone and norepinephrine (NE) concentration in the hypothalamus collected at ZT10-12 are decreased in Ud6-m-/p+ mice. Data are plotted as mean ± SEM, * p < 0.05 and ** p < 0.01 by two-tail unpaired T test. WT: N=6, Ud6-m-/p+: N=10. (B) 5-HT, DA, and Dopac contents are not significantly different in the hypothalamus of Ud6-m-/p+ mice as compared with WT mice.

### Sleep and siestas

Jones and coworkers [Jones et al., 2015] reported that Ud5-m-/p+ mice have a blunted response to sleep pressure, both during their active period and in response to forced sleep deprivation. For example, Ud5-m-/p+ mice lack the small increase in sleep late in the active phase (“siesta”) that is typical of WT mice without access to running wheels. We similarly found that Ud6-m-/p+ mice skipped the siesta near the end of the dark interval when they did not have access to running wheels. Under LD12:12 cycles, hourly assessments of wake, NREM sleep and REM sleep showed Ud6-m-/p+ mice had more wake, and less NREM sleep and REM sleep in this later part of the dark/night phase than did WT mice (Fig. 8; ZT20, p=0.017 wake; p=0.012 NREM; p=0.07 REM). Otherwise, Ud6-m-/p+ and WT mice had similar amounts of wake, NREM sleep, and REM sleep in both the dark and light periods. Ud6-m-/p+ mice also had fewer bouts of wake during the light period than WT mice (Table 1). Fast Fourier analysis of the EEG revealed Ud6-m-p+ mice have lower delta power (0.25-0.75Hz) across sleep and wake states (Suppl Figure S4). Additionally, Ud6-m-/p+ mice have increased power in the 3-4.25 Hz range in wake (Suppl Figure S4). Polyspike events with frequencies within the range of increased power in Ud6-m-/p+, and as described by Born and coworkers [Born et al., 2017], were observed throughout the EEG trace in Ud6-m-/p+ mice (Suppl Figure S4).

**Figure 8.**
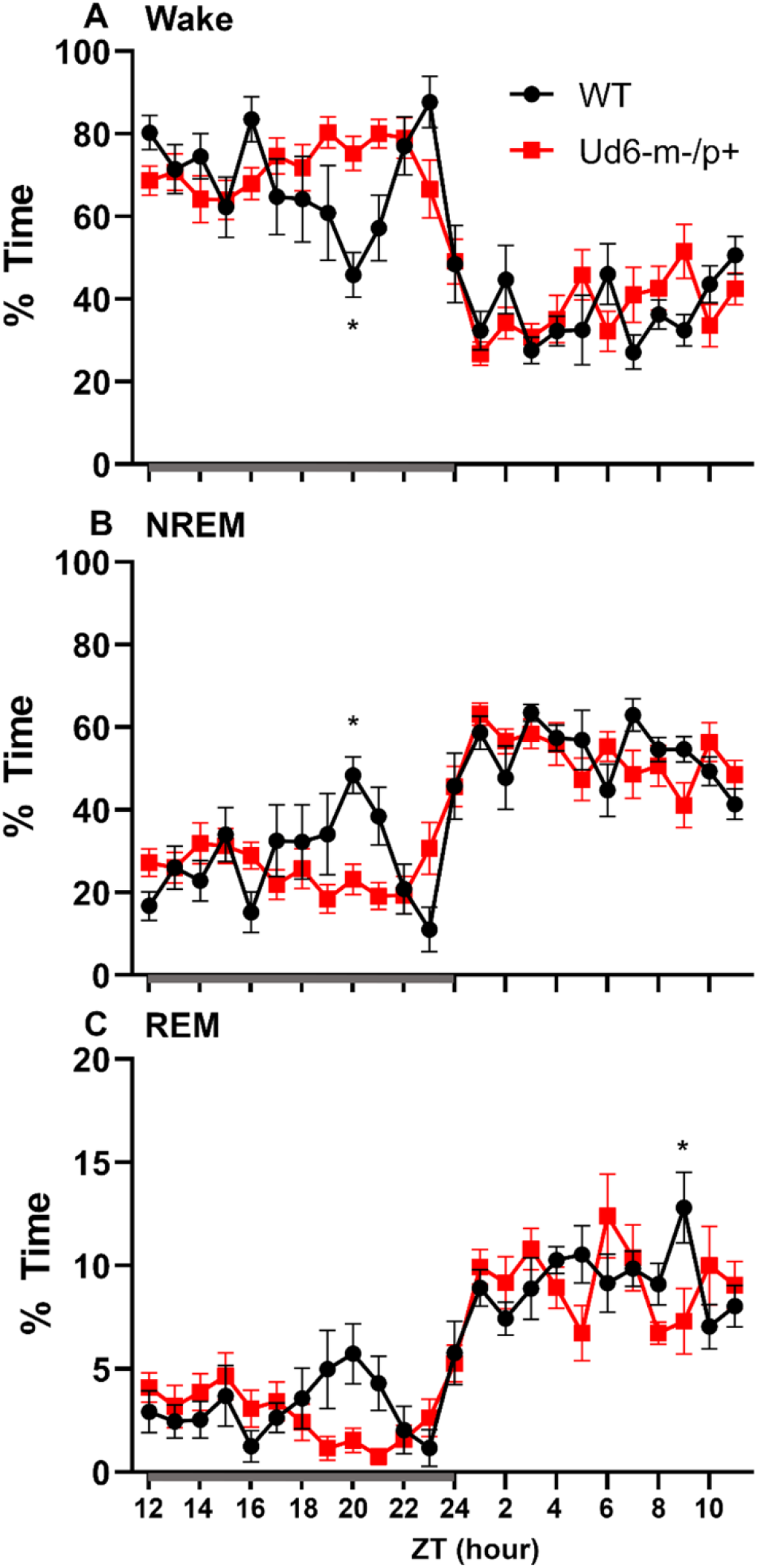
Sleep/wake behavior in WT and Ud6-m-/p+ mice in LD without access to running wheels. Amount of time in Wake (A), NREM sleep (B), and REM sleep (C) are similar in WT (black) and Ud6-m-/p+ mice (red) except that Ud6-m-/p+ mice have a blunted “siesta” episode late in the active (dark) interval. Darkness is indicated by the gray horizontal bar from ZT12-24. Data are plotted as mean ± SEM. Asterisk (*) indicates a significant difference between Ud6-m-/p+ vs. WT mice at the p<0.05 level.

**Table 1.** The number and duration of sleep and wake bouts in WT and Ud6-m-/p+ mice in LD without access to a running wheel (mean ± SEM, * p<0.05).

| **Table 1** |  | **amount (%)** |  | **# of bouts** |  | **bout duration (sec)** |  |
| --- | --- | --- | --- | --- | --- | --- | --- |
|  |  | **12hr dark** | **12hr light** | **12hr dark** | **12hr light** | **12hr dark** | **12hr light** |
| *Wake* | WT | 69.2 ± 2 | 37.9 ± 1.3 | 136.3 ± 26.5 | 176.1 ± 14.1 | 264.6 ± 38.6 | 97.1 ± 8.1 |
|  | Ud6-m-/p+ | 72 ± 2 | 38.8 ± 2 | 112.5 ± 5.8 | **141.3 ± 5.5\*** | 282.9 ± 18.6 | 124.0 ± 9.1 |
| *NREM* | WT | 27.7 ± 1.7 | 53.2 ± 1.2 | 132.9 ± 26.9 | 181.6 ± 16.7 | 107.1 ± 13.0 | 131.1 ± 7.8 |
|  | Ud6-m-/p+ | 25.3 ± 1.9 | 52.3 ± 2 | 110.8 ± 5.6 | 154.3 ± 6.3 | 106.2 ± 9.2 | 142.4 ± 6.7 |
| *REM* | WT | 3.1 ± 0.3 | 9 ± 0.2 | 29.9 ± 3.5 | 67.3 ± 5.5 | 48.8 ± 5.5 | 59.8 ± 4.1 |
|  | Ud6-m-/p+ | 2.7 ± 0.3 | 8.9 ± 0.7 | 27.3 ± 3.7 | 67.5 ± 4.1 | 46.0 ± 3.4 | 57.0 ± 2.5 |

Sleep data under LD conditions. \* vs. WT, p<0.05 unpaired T-test.

On the other hand, the propensity to siesta reverses between WT and Ud6-m-/p+ mice when the mice have access to running wheels, which tend to promote arousal, likely due to increased overall activity [Edgar et al., 1991; Espana et al., 2007]. When the mice are in LD12:12 with access to a running wheel (rotating-saucer wheel, condition “LDRW”), hourly analyses suggested that Ud6-m-/p+ mice tend to have less wake and more NREM and REM sleep in the middle of the dark period (Fig. 9A-C; WT v Ud6m-/p+ were significantly different at ZT17, p=0.045). Ud6-m-/p+ and WT mice had similar numbers of bouts and durations of each stage across the 12 h dark and light intervals (Table 2). Interestingly, Ud6-m-/p+ mice had significantly lower core body temperature than WT mice during the dark phase (Fig. 9D). Ud6-m-/p+ mice also exhibited less general activity (DSI) and running-wheel activity than WT mice during the dark phase of LD (Fig. 9E,F).

**Figure 9.**
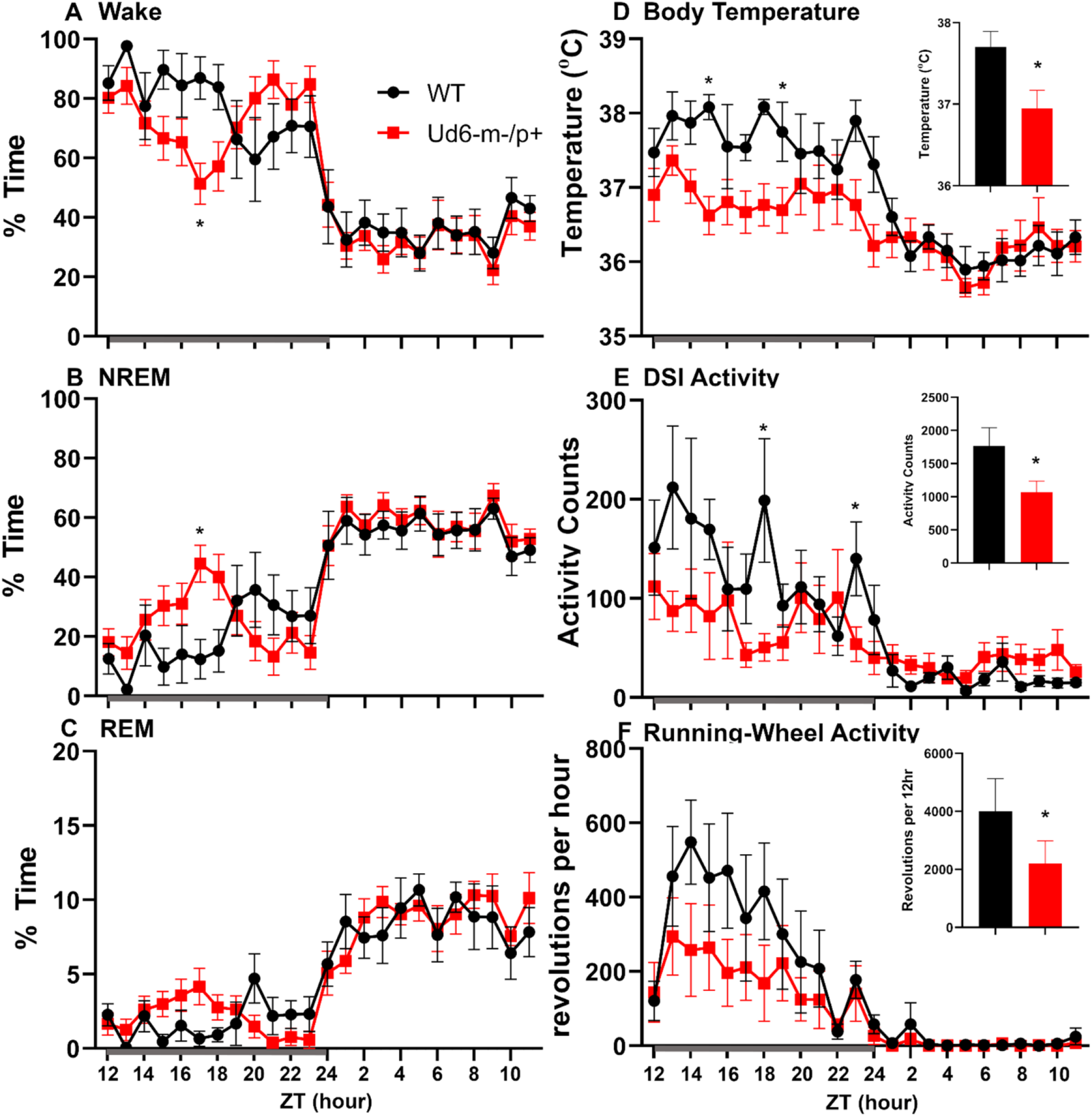
Sleep/wake behavior in LD for mice with access to running wheels. Amounts of wake (A), NREM sleep (B), and REM sleep (C) are similar in WT (black) and Ud6-m-/p+ mice (red). During the dark interval, Ud6-m-/p+ mice have lower body temperature (D), and reduced activity as recorded by DSI telemetry (E) and running-wheel revolutions (F). Darkness is indicated by the gray horizontal bar from ZT12-24. Insets summarize the data for the 12-hour dark interval for WT (black) and Ud6-m-/p+ (red). Data are plotted as mean ± SEM. Asterisk (*) indicates a significant difference between Ud6-m-/p+ vs. WT mice at the p<0.05 level.

**Table 2.** The number and duration of sleep and wake bouts in WT and Ud6-m-/p+ mice in LD with access to a running wheel (mean ± SEM).

| **Table 2** |  | **amount (%)** |  | **# of bouts** |  | **bout duration (sec)** |  |
| --- | --- | --- | --- | --- | --- | --- | --- |
|  |  | **12hr dark** | **12hr light** | **12hr dark** | **12hr light** | **12hr dark** | **12hr light** |
| *Wake* | WT | 78.3 ± 4.1 | 36.4 ± 2.4 | 73.5 ± 14.5 | 159.5 ± 33.5 | 479.9 ± 134.7 | 95.9 ± 19.9 |
|  | Ud6-m-/p+ | 73 ± 1.9 | 33.3 ± 2.3 | 67 ± 17.3 | 137.8 ± 25.0 | 711.3 ± 224.3 | 111.8 ± 16.7 |
| *NREM* | WT | 19.9 ± 3.7 | 55.3 ± 2.5 | 64 ± 18 | 163.5 ± 28.5 | 144.2 ± 0.5 | 162.2 ± 33.6 |
|  | Ud6-m-/p+ | 24.9 ± 1.8 | 58 ± 2.2 | 59.4 ± 17.5 | 151.8 ± 19.6 | 146.5 ± 17.1 | 187.9 ± 47.3 |
| *REM* | WT | 3.3 ± 1 | 9.4 ± 1.2 | 12 ± 1 | 47.5 ± 20.5 | 55.9 ± 30.5 | 64.1 ± 2.3 |
|  | Ud6-m-/p+ | 2.1 ± 0.2 | 8.6 ± 0.3 | 10.2 ± 4.2 | 65.2 ± 7.6 | 42.8 ± 9.5 | 62.4 ± 10.4 |

Sleep data under LDRW conditions.

Under constant dark conditions with access to a running wheel (condition “DDRW”), Ud6-m-/p+ mice had less wheel running than WT mice in the first part of the active phase, and their body temperature was correspondingly lower. Ud6-m-/p+ mice also had less wake and more NREM and REM sleep over this initial part of the active phase than WT mice (Suppl Figure S5). Neither WT nor Ud6m-p+ mice had a siesta under the DDRW condition. Ud6-m-/p+ mice had less wake (v. WT p=0.03) and more NREM sleep (p=0.042) at CT14. Ud6-m-/p+ mice also spent less time and had shorter REM sleep bout duration than WT mice during the subjective light (Supplemental Table 1). Overall, the lower amounts of wake in Ud6-m-/p+ mice during the early subjective dark period may be related to lower locomotor activity and body temperature.

### Sleep deprivation

Jones and coworkers [Jones et al., 2015] also reported that Ud5-m-/p+ mice have a reduced response to sleep deprivation. Typically, long periods of wake are followed by high EEG delta power in the subsequent NREM sleep. We found that after 6-h of sleep deprivation at the start of the light, Ud6-m-/p+ mice had lower delta power (0.25 to 4 Hz) in NREM sleep during the first hour of recovery sleep compared to WT mice (Suppl Figure S6), though delta power then followed a normal pattern in the subsequent hours. Other measures of increased sleep pressure, including amount of time, number of bouts and duration of bouts of each state (wake, NREM sleep and REM sleep) during recovery were similar between Ud6-m-/p+ and WT mice (Suppl Table 2).

## Discussion

The new Ud6-m/p+ model has the potential to become a preferred rodent system for understanding AS. It combines the advantages of mouse genetics over the rat model of AS [Dodge et al., 2020] with the likelihood of enhanced stability and consistency over the Ud5-m-/p+ model because of its 4X larger deletion (1247 nt). Moreover, Ud6-m-/p+ specifically targets the *Ube3a* gene so that its phenotypes can be attributed to that single gene, whereas a liability of the UG-m-/p+ model mouse is that two other genes are also affected, which complicates interpretations [Jiang et al., 2010]. We confirmed that the Ud6-m-/p+ mouse exhibits standard phenotypes observed in the other rodent models of AS [Dodge et al., 2020; Jiang et al.,1998; Jiang et al., 2010; Sonzogni et al., 2018], such as worse coordination on rotarod and reduced marble burying (Fig. 4).

In contrast to previous reports on the Ud5-m-/p+ mouse model [Ehlen et al., 2015; Jones et al., 2016], we clearly find much lower UBE3A protein expression in the SCN of Ud6-m-/p+ mice, thereby vindicating the expectation that *Ube3a* is imprinted in neurons, and that SCN neurons are not exceptional in that regard. These differing conclusions may be due to the aforementioned tendency of the Ud5-m-/p+ model towards strain dependency and loss of phenotype over time [Born et al., 2017; Huang et al., 2013]. Another source of the discrepancy in apparent expression of UBE3A in the SCN may be due to different methods. The study of Jones and coworkers [Jones et al., 2016] used the NeuN marker to identify neurons, but the accuracy of this particular marker for neurons in the SCN has been questioned by Morin and coworkers [Morin et al., 2011]. Apparently NeuN-positive neurons represent only a subpopulation of neurons with a specific spatial distribution in the mouse SCN [Morin et al., 2011], and therefore the use of NeuN may overlook a large population of SCN neurons. In contrast, our IHC utilized an antibody to the neuron-specific RNA-binding protein, ELAVL4/HuD that has been used successfully as a global marker of neuronal cells in brain tissues [Jung & Lee., 2021]. Finally, our assessment of UBE3A expression in the SCN was based on the more stringent criterion of “number of UBE3A-positive cells,” and we found that 25% of SCN cells retain a low level of UBE3A expression (Fig. 3C). Why? Recent research may provide an explanation for this heterogeneity within the SCN as approximately 20-30% of SCN cells are neither neurons nor astrocytes [Morris et al., 2021; Wen et al., 2000; Xu et al., 2021]. Using the ELAVL4 antibody to define which cells are neuronal vs. non-neuronal [Jung & Lee., 2021], we found that SCN neurons of the Ud6-m-/p+ mouse have significantly reduced expression of UBE3A [Supplementary Fig. S1]. Moreover, it seems likely that some of the UBE3A-positive cells are not neurons or astrocytes, but represent oligodendrocytes, endothelial or ependymal cell types that do not paternally imprint *Ube3a* [Grier et al., 2015; Jiang et al., 2010; Yamasaki et al., 2003]. Therefore, our conclusion is that UBE3A expression is dramatically suppressed in SCN neurons of Ud6-m/p+ mice, and therefore the paternal *Ube3a* gene is imprinted in SCN neurons as expected from studies of neurons in other brain regions [Albrecht et al., 1997; Jiang et al., 1998; Weeber et al., 2003].

We previously reported that circadian robustness was impaired in the Ud5-m-/p+ and UG-m-/p+ mouse models of AS [Shi et al., 2015]. Here, we show that the Ud6-m-/p+ model shares these same phenotypes, especially the suppressed locomotor activity in LL (Fig. 5). We also observed depressed locomotor activity rhythms (both running wheel and total DSI activity) in LD and DD (Fig. 9 & Supplementary Fig. S4). Consequently, we observe significant circadian phenotypes in all three mouse models of Angelman syndrome, and our results concur with an earlier circadian study of the roles of *Ube3a* in tissue cultures [Gossan 2014]. Inexplicably, Ehlen and coworkers [Ehlen’2015] could not repeat our prior observation of lengthened circadian periods in Ud5-m-/p+ mice (although their data showed a trend towards lengthening, the difference was not statistically significant). We attribute these differences to unidentified differences in the conditions under which the rhythms were monitored in the various labs, or to the lability of the Ud5-m-/p+ model, including a potential loss of phenotype over time that might have affected the mice in other labs [Born et al., 2017; Huang et al., 2013; Dodge et al., 2020]. Our observations here with the Ud6-m-/p+ mouse model reaffirms that *Ube3a* deficiency has circadian ramifications. The present study also demonstrates reciprocal effects of *Ube3a* gene dosage on the circadian system. When UBE3A levels are reduced in neurons due to paternal imprinting, light suppresses wheel-running activity and the circadian period lengthens (Fig. 5 and [Shi et al., 2015]). Conversely, light enhances the activity when mice are in the light, and the circadian period of the cultured SCN clock *in vitro* is shorter when Ube3a is over-expressed in an autism mouse model (Fig. 6). Therefore, the circadian clock is inversely sensitive to loss and gain of *Ube3a* expression, supporting an important homeostatic action for UBE3A in the circadian clockwork.

Children with Angelman syndrome often have nighttime awakenings and difficulty initiating sleep [Heal and Oliver et al., 2017; Tricket et al., 2017]. In mouse models of Angelman syndrome, two previous studies reported altered sleep patterns in an Angelman syndrome mouse line [Ehlen et al., 2015; Copping et al., 2021]. Ehlen and co-workers [Ehlen et al., 2015] reported a lack of the “siesta-like” reduction of wake in the late night/active bout of Ud5-m-/p+ mice compared to WT mice. Similarly, under LD conditions in our experiments, Ud6 m-/p+ mice did not show this siesta which was clear in WT mice around ZT20 (Figure 8). Interestingly, when a running wheel is present, the Ud6-m-/p+ mice are active later in the dark/night interval (Figure 9A), which is a phased activity pattern reminiscent to that we reported for Ud5-m-/p+ mice [Shi et al., 2015] and which we attributed to the entrainment of a longer free-running period to LD. Still, even with access to a running wheel, Ud6-m-/p+ mice had lower core body temperature, most likely due to less wheel running than their WT littermates (Figure 9D/E/F).

Increased EEG delta power during wake and sleep has been reported in children with AS and a mouse model of AS [Bakker et al., 2018; Sidorov et al., 2017,], whereas lower delta power late in the dark period was reported for the Ud5-m-/p+ mouse model by Ehlen and co-workers [Ehlen et al., 2015]. The increased high delta/low theta power observed in the Ud6-m-/p+ mice is similar to previous reports of altered power distribution in the EEG of *Ube3a* deficient mice. Born and colleagues [Born et al., 2017] reported increased theta power in *Ube3a* maternal deficient mice and attributed this increase to polyspike events in cortical and hippocampal EEG traces. We also observed polyspike events in our cortical EEG recordings mainly in wake and REM sleep, with fewer in NREM sleep. These events may indicate a predisposition for seizure activity, but we did not observe overt seizures in these mice [Born et al., 2017]. Unfortunately, Born and co-researchers did not separate the power spectrum analysis by state of arousal and so comparisons between studies are difficult. The low delta power observed in the Ud6-m-/p+ mice differ from other groups, however the differences in scoring stages of wake and sleep may account for some of the discrepancy, in addition to different genetics of the AS models. Moreover, Jones and coworkers reported that Ud5-m-/p+ mice have a reduced capacity to accumulate sleep pressure in response to sleep deprivation [Jones et al., 2015]; typically, increasing duration of wake increases EEG delta power in the subsequent NREM sleep. We found that Ud6-m-/p+ mice had lower delta power during their initial NREM sleep after 6-h of sleep deprivation, suggesting some blunting of the response to sleep pressure.

Ud6-m-/p+ mice also differed in their pattern of REM sleep. We found that Ud6m-/p+ had ∼30% less REM sleep amount during the subjective day of DD (with access to a running-wheel), most likely due to shorter bouts of REM sleep. Similarly, a prior study reported that Ud5-m-/p+ mice had about 20% less REM sleep across the 24-h day [Ehlen et al., 2015]. Though their scoring methods differ, Copping & Silverman [Copping & Silverman., 2021] also report less REM sleep in Ube3a null mice. Future research in people with AS might target whether their fragmented sleep disrupts REM sleep.

Pharmacological and genetic studies in mammals suggest that norepinephrine plays a critical role in promoting arousal, especially during conditions that require high attention or activation of the sympathetic nervous system [España et al., 2011]. Moreover, corticosteroid therapy benefits children with Angelman syndrome [Forrest et al., 2009]. We detected reduced norepinephrine in the hypothalamus of Ud6-m-/p+ mice during the late day (ZT10-12 in Fig. 7), which correlates with a decreased wake during the subsequent night (ZT17 in Fig. 9). Reduced levels of norepinephrine during the early subjective night (CT14 of DD) in Ud5-m-/p+ cerebellum is also in line with decreased wakefulness during early night in Ud6m-/p+ animals (Supplementary Figures S3 & S4). Low norepinephrine is also associated with drowsy and inattentive behaviors in mammals [España et al., 2011], which correlates with symptoms of Angelman patients [Pelc et al., 2008]. Stress hormones, including corticosterone and norepinephrine, are released during and immediately after stressful stimuli tested in emotionally arousing learning tasks [Aston-Jones et al., 2005]. Moreover, removal of these hormones by adrenalectomy impairs memory consolidation for emotionally arousing experiences [Foote et al., 1980; Roozendaal et al., 2002]. Observations from human AS patients correlate well with our measurements indicating depleted levels of corticosterone and norepinephrine in *Ube3a* deficient mice [Kim et al., 2018], and may represent a partial explanation for compromised cognition and impaired learning & memory in AS. Taken together, the hypothalamic-pituitary-adrenal (HPA) axis and may represent new therapeutic targets to ameliorate AS phenotypes. Moreover, cortisol and norepinephrine may serve as easily measurable biomarkers for assessing the efficacy of potential therapeutic treatments of Angelman syndrome in clinical trials.

Although animal models can never fully recapitulate the complexity of clinically important human syndromes, they can provide key resources for analyses. In this case, mouse models of Angelman syndrome help to identify the mechanistic consequences of alterations of *Ube3a* expression levels that result in Angelman syndrome when deficient versus autism when over-abundant. The impact of *Ube3a* under- and over-expression is considerable upon human neurodevelopment and sleep, and the new Ud6-m-/p+ mouse provides an experimentally tractable model to analyze sleep and the underlying regulation by the circadian system.

## Materials and methods

### Mouse strains

C57BL/6J mice harboring a knockout of exon 6 of the *Ube3a* gene were generated with CRISPR/Cas9 (Fig. 1A), and this strain is herein called “Ud6-m-/p+.” We confirmed that the Ud6-m-/p+ strain in our mouse colonies was genetically 98.97% (289/292 SNPs) C57BL/6J (JAX# 000664), as assessed by genotyping performed by the Jackson Laboratory (Bar Harbor, ME). Female mice harboring one *Ube3a* knockout allele were crossed with wild-type background (WT C57BL/6J) male mice to obtain the maternal deletion mice (Ud6-m-/p+) that are the model for Angelman syndrome. “Ud5-m-/p+” represents the original Angelman model mouse strain harboring an exon 5 deletion in *Ube3a* (Jiang et al., 1998).

*Ube3a* conventional BAC overexpressing mouse models (*Ube3a*-Tg 1x) on the FVB background [Krishnan et al., 17; Smith et al., 2011] were crossed with C57Bl6 WT or Per2::Luc mice to generate an F1 generation with a mixed genetic background) to obtain the WT, *Ube3a*-Tg 1x, and *Ube3a*-Tg 2x strains shown in Figure 6. As reported by other research groups for “Angelman phenotypes” [Born et al., 2017; Huang et al., 2013], we also found that mouse genetic backgrounds have significant effects on circadian properties, e.g., the period of the SCN rhythm *in vitro* displayed a lengthened circadian period on the FVB/C57BL/6 mixed background (Fig. 6A) compared with that on the C57BL/6 background [Shi et al., 2015].

Age and sex balanced experimental and littermate control mice were used in all animal assays. At Vanderbilt University, mice were housed in temperature-controlled environments on LD12:12 (12:12-h light-dark cycle with lights-on at 06:00, cool-white fluorescence light ∼ 300 lux) and fed chow (5001, Lab diet, 5% fat) for breeding and experiments. All animal experiments were approved by the Vanderbilt University and Beth Israel Deaconess Medical Center Institutional Animal Care and Use Committees and were conducted according to those committees’ guidelines.

### Immunoblotting and quantitative RT-PCR

Animals were sacrificed by cervical dislocation at ZT4 in LD 12:12. The cerebellum samples were collected and stored at -80°C until protein and RNA were extracted. Tissues were sonicated in RIPA buffer containing a cocktail of protease inhibitors (Sigma) and clarified by centrifugation (RIPA buffer = 50mM Tris pH 8.0, 150 mM NaCI, 0.1% SDS,1.0% NP-400, 5% Sodium Deoxycholate). Protein from the supernatant was quantified by the Bradford assay (Bio-Rad). Proteins were separated by SDS-PAGE and then transferred to PVDF membranes for immunoblotting. Monoclonal IgG directed against UBE3A protein (E8655, Sigma) and anti-β-ACTIN monoclonal antibody (A5316, Sigma) were used to probe UBE3A and β-ACTIN, respectively. To test the UBE3A and FLAG tag transgenic expression in the SCN, protein extracts from SCN cultures were examined by SDS-PAGE immunoblotted with anti-UBE3A monoclonal antibody (E8655, Sigma), or anti-FLAG monoclonal antibody (F1804, Sigma). Densitometric analyses were performed using Image J software (NIH).

Total cerebellum RNA collected at ZT4 were extracted and transcribed by Superscript III reverse transcriptase (Invitrogen) following the manufacturer’s instructions. Diluted cDNA was amplified by PCR using the Sybr Green Master Mix (Sigma), and raw threshold-cycle time (Ct) values were calculated with iQ5 software (Bio-Rad). Mean values were calculated from samples and normalized to those obtained for *Hprt* transcripts as reported previously [Shi et al., 2015].

### Marble burying and Rotarod assay

In order to assess anxiety & compulsiveness, Ud6-m-/p+ mice were placed in individual cages containing ∼5 cm of diamond soft bedding (Teklad 7089C) at ZT 3-5 for 15 minutes to acclimate to the test conditions, which were in the light (mice in subjective daytime). After 15 minutes, each mouse was briefly removed from its cage, and twenty-five marbles (∼1.5 cm) were placed in the cage in 5 rows. The mice were returned to the cage and allowed 30 minutes to investigate the marbles. The number of marbles buried at least two-thirds of the way in the bedding was recorded. To test the anxiety during the night phase, mice from their home cage at ZT 15-17 were transferred to a testing cage in the dark under infrared illumination. The mice were tested in the dark (mice in subjective nighttime) in the test cage with marbles in diamond soft bedding. After 30 minutes, mice were returned to their home cages in the dark under infrared illumination, and the number of marbles buried in the bedding is recorded.

The ability to maintain balance on a rotating cylinder is measured with a standard rotarod apparatus (Ugo Basile, Gemonio, Italy), consisting of a cylinder (approx. 3 cm in diameter) that rotates at speeds ranging from 4 to 40 rpm in 5 minutes. Mice are placed on the rotarod and confined to a section of the cylinder approximately 6.0 cm wide by Plexiglas dividers. The latency at which mice fall off the rotating cylinder is automatically measured. Rotarod performance was assessed three times daily during ZT 8-11 for two contiguous days.

### Per2:Luc organotypic slice bioluminescence assays

Mice harboring the P_mPer2_::mPER2-LUC knock-in allele [Yoo et al., 2004] that have been backcrossed with the C57BL/6J strain for more than 12 generations were crossed with the *Ube3a* BAC transgenic mice (FVB background) to obtain *Ube3a*-Tg 1x with the luminescence reporter of mPER2 expression. The data in Fig. 6 were measured from these mixed background mice in the F1 generation so that they were 50% FVB and 50% C57BL/6J. One to two hours before lights-off of LD 12:12, the mice were sacrificed by cervical dislocation, and cultures of SCN and peripheral tissues were prepared as previously described [Shi et al., 2015]. The tissues were cultured in recording medium containing 10% fetal bovine serum (Gibco) and 100 nM of luciferin (Promega) at 36.5°C. The *in vitro* rhythms were analyzed using LumiCycle analysis software (Actimetrics, Evanston, IL).

### Determination of biogenic amine and corticosterone concentration

The Ud6-m-/p+ and WT hypothalamus and serum samples were collected from mice sacrificed at ZT10-12 in LD 12:12. The Ud5-m-/p+ and WT cerebellum samples were collected from mice in DD for 26 h and 38 h. Biogenic monoamines in hypothalamus and cerebellum were analyzed using LC/MS by the Neurochemistry Core Laboratory of Vanderbilt University Medical Center, and the samples were normalized to total protein concentration. The corticosterone concentration in serum was analyzed using double antibody RIA by the Hormone Assay and Analytics Core Laboratory of Vanderbilt University Medical Center.

### Immunohistochemistry (IHC)

The animals were deeply anesthetized by Avertin (1.25%, Sigma) and perfused in PFA (4% in PBS, Sigma). Brains were removed and postfixed in 4% PFA for 12 h at 4 °C, and cryoprotected in 20% sucrose-PBS overnight at 4 °C. Coronal 30-μm-thick sections were cut on a freezing microtome (CM1850, Leica, Germany) and free-floating sections were processed for standard avidin-biotin IHC and immunofluorescence methods as previously described [Liska et al., 2021; Sumová et al., 2002]. The sections were incubated in 0.5% peroxide solution in PBS (0.01M sodium phosphate-0.15 M NaCl, pH 7.2) for 10 min for peroxidase blocking. For the immunofluorescence, this step was skipped. Then, the sections were incubated in 2% normal goat serum (Vector Laboratories, USA) + 1% bovine serum albumin (Sigma, USA) + 0.3% Triton X-100 (Sigma, USA) for 60 min.

The primary polyclonal antibodies applied overnight at 4 °C were as follows: mouse anti-UBE3A 1: 600 (SAB1404508, Sigma, USA); rabbit anti-ELAVL4/HudD 1: 600 (ab96474, Abcam, Great Britain) as a neuronal marker [Loffreda et al., 2020; Okano et al., 199,]; chicken anti-GFAP 1: 1000 (ab4674, Abcam, Great Britain) as a brain astrocyte marker [Brenner et al., 1994]. For the immunofluorescence IHC, the secondary antibodies (1: 600; 1 h) were goat anti-mouse fluorescent (F2761, ThermoFischer Scientific, USA), goat anti-rabbit fluorescent (A11037, ThermoFischer Scientific, USA), and goat anti-chicken fluorescent (ab97145, Abcam, Great Britain). The immunofluorescence was visualized using confocal microscopy (Multiphoton Microscope Leica TCS SP8 MP, Leica, Germany).

For the avidin-biotin IHC, we applied standard methods with diaminobenzidine tetrahydrochloride (Sigma, USA) as the chromogen (Vector Laboratories, Peterborough, UK). For quantification, cell nuclei labeled irrespective of the intensity of their staining were counted in the whole SCN (3-9 sections per brain) by a “procedure-blind” person using an image analysis system (ImageJ/FUJI). In the hippocampal regions, where cell bodies are dense and difficult to reliably resolve, optical staining density was measured (3-5 sections per brain) rather than counting individual cells for positive staining. After conversion of the images into grey scale, the optical density measurement was performed selectively in CA1, CA3 and DG regions and in the closely adjacent areas without cell bodies. The optical density (grey color intensity) was calculated as follows: OD = log (background intensity / cells intensity). For the density measurements, the sections from WT and Ud6-m-/p+ mice were assayed simultaneously in one IHC assay.

### Sleep EEG procedures

For sleep EEG measurements at the Beth Israel Deaconess Medical Center, male and female mice were housed on LD12:12 (lights on at 07:00) with constant temperature (22 ± 1.4°C) and humidity (26 ± 2.1 mmHg). Regular chow (LabDiet #5058) and water were available *ad libitum*. Mice were 16.4 ± 0.9 weeks old and weighed 25.4 ± 0.5 g at the time of surgery. We anesthetized mice with ketamine/xylazine (100/10 mg/kg, i.p.) and placed them in a stereotaxic alignment system (model 1900; Kopf Instruments). Electroencephalogram (EEG) and electromyogram (EMG) electrodes were soldered to a 2 × 2 pin microstrip connector and implanted and affixed to the skull with dental cement for polysomnogram recordings. For the EEG electrodes, two stainless steel screws were implanted into the skull 1.5 mm lateral and 0.5 mm rostral to bregma and 1.0 mm rostral to lambda. The EMG electrodes were made from multistranded stainless steel wire (Cooner Wire) and placed into the neck extensor muscles. We treated each mouse with meloxicam (4 mg/kg slow release, s.c.) at the time of surgery.

### Sleep recordings

One week after implantation of EEG and EMG electrodes, we transferred mice to recording cages in a sound-attenuated box. We attached the recording cable to a low-torque electrical swivel that was fixed to the cage top to allow free movement. We allowed the mice 7 days to habituate to the recording cable and chamber. EEG/EMG signals were amplified, filtered (high-pass, 0.3 Hz; low-pass, 1 kHz), digitized at a sampling rate of 256 Hz, and recorded with SleepSign software (Kissei Comtec). We used SleepSign (filter settings: EEG, 0.25–64 Hz; EMG, 10–60 Hz) for preliminary, semiautomatic scoring of wake, REM, and non-REM (NREM) sleep in 10 s epochs and then examined all epochs and made corrections when necessary.

We recorded sleep/wake behavior under 4 conditions for WT (n=8) and Ud6m-/p+ (n=12). Twenty-four-hour sleep recordings were first acquired under LD12:12 conditions, after which time the response of the mice to sleep deprivation was assessed. At the start of the light phase, mice were kept awake for 6 h by introducing novel objects into the cage or by lightly tapping on the side of the cage, and subsequently behavior across the 6-h recovery period after the sleep deprivation was recorded. Then, mice were acclimated to running wheels for 10 d and 24-h sleep recordings were acquired in the “LD with running wheel” (LDRW) condition. Finally, mice were held in constant dark (DD) for at least 14 d prior to performing 24-h sleep recordings under “DD with running wheel” (DDRW) condition. RW, LMA and Tb data were analyzed using Clocklab (Coulbourne Instruments, Natick, MA) to analyze period (Chi-squared periodogram) and cosinor amplitude for the last 7 days in LD and the last 7 days in DDRW. Two-way repeated measures ANOVA was used to compare between genotypes with Bonferroni tests for multiple comparisons serving as post hoc analyses. For DDRW sleep data plots, running wheel onset were manually determined to align the start of subjective dark.

### Circadian locomotor behavior measurement

For the circadian behavioral experiments performed at Vanderbilt University, age- and gender-matched littermates were singly housed in cages equipped with running-wheel or infrared sensors for locomotor activity measurements with unlimited access to regular chow (5001, Lab diet) and water. For the running wheel experiments shown in Figure 8, a rotating-saucer running-wheel was used to measure the locomotor activity at Beth Israel Deaconess Medical Center. ClockLab software (Actimetrics, Evanston, IL) was used to collect data and perform period and activity analyses. Circadian periods in DD and LL were calculated using Chi-Square periodogram analyses.

### Statistical Analyses

Data are presented as means ± SEM. Statistical analyses were performed by two-tail unpaired t-test, one-way, and two-way ANOVA as indicated (GraphPad, PRISM9). For sleep EEG, we analyzed light and dark period data between mutants and WT littermates using unpaired T-tests. Hourly data was assessed with two-way ANOVA (time x genotype) and post hoc Bonferroni tests were run to adjust for multiple comparisons (GraphPad, PRISM9). Recovery sleep NREM spectra was analyzed with area under the curve including frequencies of 0.25-4Hz.

## Abbreviations

LD: light/dark cycle (LD 12:12 = 12-h light/12-h dark cycle)
LL: constant light conditions
DD: constant darkness conditions
CT: circadian time, where CT0 is subjective dawn and CT12 is subjective dusk
ZT: phase/time in the LD cycle, where ZT0 is lights-on and ZT12 is lights-off in a LD 12:12 cycle

## Acknowledgements

We thank N. Machado and W. Todd for technical assistance with the sleep studies, and Ms. Dan Cai for assistance with breeding Angelman model mice. The Vanderbilt Murine Neurobehavior Core lab (U54HD083211, NICHD) assisted in the performance of the Rotarod and Marble Burying testing, the Vanderbilt Neurochemistry Core Lab assayed biogenic amines, and the Vanderbilt Hormone Assay and Analytics Core Laboratory (DK059637 & DK020593, NIDDK) determined corticosterone levels. Dr. Sumová thanks MEYS (LM2015062 Czech-BioImaging) for support. This work was supported by funding from the USA National Institutes of Health (R01 NS104497 to CHJ & TES, R01 NS106032 to TES, and R01MH112714 & R01MH114858 to MPA). We are grateful to Dr. Terri Jo Bichell for inspiring us to study circadian rhythms and sleep in Angelman models.

## Supplemental Figures

**Supplemental Figure 1. Paternal *Ube3a* gene is imprinted in neuronal but not glial cells of the SCN and hippocampus of Ud6-m-/p+ mice**.

(A) High magnification confocal images of quadruple immunofluorescence for UBE3A, glial fibrillary acidic protein (GFAP, a marker for glial cells), ELAV-like RNA Binding Protein 4 (ELAVL4, a marker for neurons), and diamidino-2-phenylindole (DAPI, a marker for nuclei) in the SCN of WT (upper panels) and Ud6-m-/p+ mice (lower panels). **Left panels**: Merged images of GFAP (yellow/orange), ELAVL4 (red), and DAPI (blue); arrows indicate GFAP-immunopositive cells. **Right panels**: UBE3A (green); arrows indicate absence of UBE3A staining in GFAP positive cells depicted by arrows in the left panels.

(B) Confocal images of quadruple immunofluorescence of GFAP, ELAVL4, UBE3A, and DAPI in the hippocampus WT (upper panels) and Ud6-m-/p+ mice (lower panels). **Left panels**: Merged images of GFAP (yellow/orange), ELAVL4 (red), and DAPI (blue); arrows indicate GFAP-immunopositive stained cells. **Right panels**: UBE3A (green); arrows indicate absence of UBE3A staining in the GFAP positive cells depicted by arrows in the left panels.

**Supplemental Figure 2. Sex differences in FRP of running-wheel rhythms of Angelman male mice in constant darkness (DD)**.

The free-running periods (FRPs) of wheel-running behavior on rotating saucer exercise wheels were analyzed by ClockLab software. Mice were initially housed in LD 12:12 or in DD with or without rotating saucer running-wheels, and were then transferred to DD with rotating saucer running-wheels. **Left**: Pooled data for both male and female mice (WT: n=7, Ud6-m-/p+: n=7, UG-m-/p+: n=7). **Middle**: Male mice (WT: n=4, Ud6-m-/p+: n=4, UG-m-/p+: n=5). **Right**: Female mice (WT: n=3, Ud6-m-/p+: n=3, UG-m-/p+: n=2). Data points with Mean ± SEM of the group of pooled male and female mice (**left**), and male mice (**middle**) are plotted, *p<0.05, *** p<0.001, or as indicated by two-tail unpaired T test. Data points with Mean of the group of female mice (**right**) are plotted, and no significance is detected among genotypes in female mice.

**Supplemental Figure 3. Levels of norepinephrine and other neurotransmitters in cerebellum of Ud5-m-/p+ mice**.

Mouse cerebellums were collected at CT 14 in DD (26 h in DD after release from LD 12:12). GABA and monoamine contents were analyzed by a LC/MS assay. Data are represented as the mean ± SEM (WT: N=5, Ud5-m-/p+: N=4). NE: norepinephrine, GABA: gamma-Aminobutyric acid, 5-HT: serotonin, DA: dopamine, Dopac: dihydroxyphenylacetic acid. ** p < 0.01 by two-tail unpaired T test.

**Supplemental Figure 4. Spectral analysis of Wake, NREM sleep and REM sleep in LD condition**.

Ud6-m-/p+ (red) have lower delta (0.25-0.75Hz) power than WT in both the dark and light phase during (A,A’) Wake, (B,B’) NREM sleep and (C,C’) REM sleep. Ud6-m-/p+ have more power in the range of high delta/low theta (3-4.25Hz). Polyspike episodes in this frequency were observed in Ud6-m-/p+ mice and insets of representative traces from each state are shown. Horizontal black lines indicate significance versus WT (p<0.05) following 2-way ANOVA and Bonferroni correction.

**Supplemental Figure 5. Constant dark conditions with access to a running wheel (DDRW)**. Under DDRW, amounts of wake (A), NREM sleep (B), and REM sleep (C) are similar in WT (black) versus Ud6-m-/p+ mice (red). During the initial active phase, Ud6-m-/p+ mice have lower body temperature (D), and reduced activity as recorded by DSI telemetry (E) and running-wheel revolutions (F). Active phase is indicated by the gray horizontal bar from CT12-24. Insets summarize the data for the 12-hour dark interval for WT (black) and Ud6-m-/p+ (red). Data are plotted as mean ± SEM. Asterisk (*) indicates a significant difference between Ud6-m-/p+ vs. WT mice at the p<0.05 level.

**Supplemental Figure 6. Spectral analysis of recovery sleep and wake following 6 hours of sleep deprivation**. Ud6-m-/p+ (red) have lower NREM delta power than WT (black) in the first hour of sleep recovery following 6 hours of sleep deprivation. The area under the curve was calculated for each mouse with normalized power spectrum data in the range of 0.25 to 4 Hz (* vs. WT, p<0.05).

**Supplemental Table 1. 12-hr summary of sleep under constant dark with access to a running wheel condition (DDRW)**. Data are means ± SEM. Asterisk (*) indicates a significant difference between Ud6-m-/p+ vs. WT mice at the p<0.05 level.

**Supplemental Table 2. 6-hr summary of sleep following sleep deprivation**. Ud6-m-/p+ vs. WT mice had similar responses to 6 hours of sleep deprivation. Data are means ± SEM. /p+ Sleep data under LD conditions. * vs. WT, p<0.05 unpaired T-test. Sleep data under LDRW conditions.

