## Supplement for "Altered circadian rhythms and sleep in a new Angelman Syndrome mouse model"

### Supplemental Information for Shi et al., "Altered circadian rhythms and sleep in a new Angelman Syndrome mouse model "

#### 1. Supplemental Data; Figures S1-S6

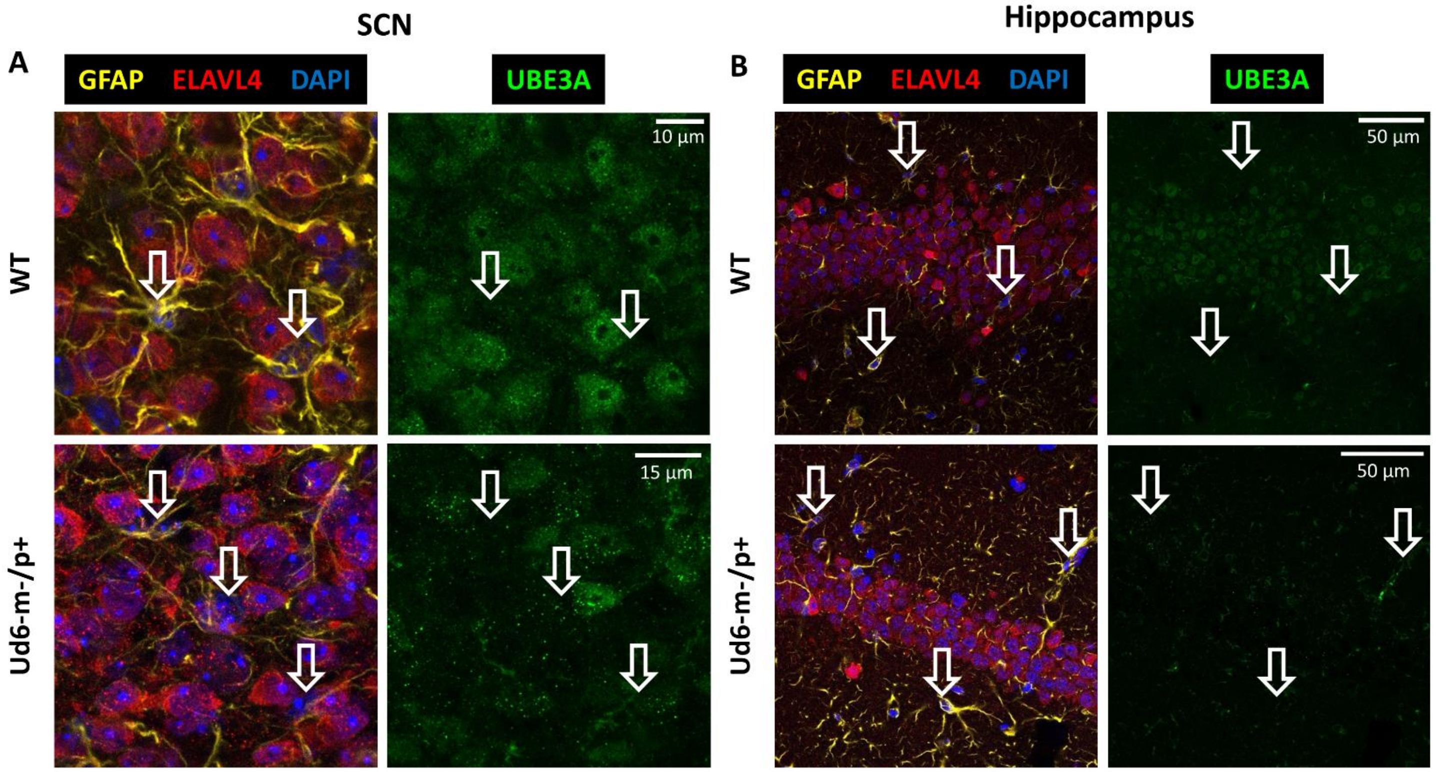


**Supplemental Figure 1. Paternal *Ube3a* gene is imprinted in neuronal but not glial cells of the SCN and hippocampus of Ud6-m-/p+ mice.**

(A) High magnification confocal images of quadruple immunofluorescence for UBE3A, glial fibrillary acidic protein (GFAP, a marker for glial cells), ELAV-like RNA Binding Protein 4 (ELAVL4, a marker for neurons), and diamidino-2-phenylindole (DAPI, a marker for nuclei) in the SCN of WT (upper panels) and Ud6-m-/p+ mice (lower panels). **Left panels**: Merged images of GFAP (yellow/orange), ELAVL4 (red), and DAPI (blue); arrows indicate GFAP-immunopositive cells. **Right panels**: UBE3A (green); arrows indicate absence of UBE3A staining in GFAP positive cells depicted by arrows in the left panels.

(B) Confocal images of quadruple immunofluorescence of GFAP, ELAVL4, UBE3A, and DAPI in the hippocampus WT (upper panels) and Ud6-m-/p+ mice (lower panels). **Left panels**: Merged images of GFAP (yellow/orange), ELAVL4 (red), and DAPI (blue); arrows indicate GFAP-immunopositive stained cells. **Right panels**: UBE3A (green); arrows indicate absence of UBE3A staining in the GFAP positive cells depicted by arrows in the left panels.

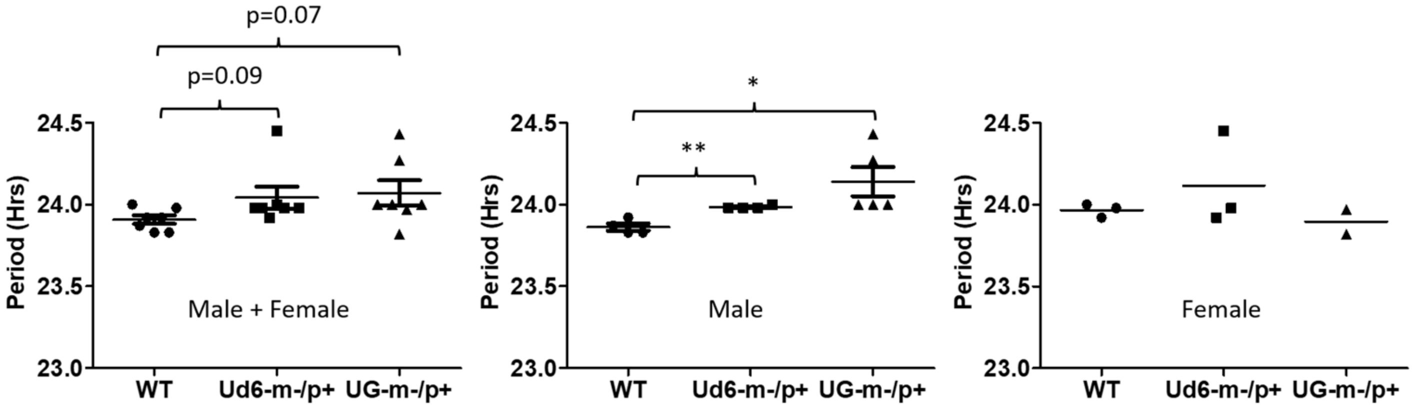


**Supplemental Figure 2. Sex differences in FRP of running-wheel rhythms of Angelman male mice in constant darkness (DD).**

The free-running periods (FRPs) of wheel-running behavior on rotating saucer exercise wheels were analyzed by ClockLab software. Mice were initially housed in LD 12:12 or in DD with or without rotating saucer running-wheels, and were then transferred to DD with rotating saucer running-wheels. **Left**: Pooled data for both male and female mice (WT: n=7, Ud6-m-/p+: n=7, UG-m-/p+: n=7). **Middle**: Male mice (WT: n=4, Ud6-m-/p+: n=4, UG-m-/p+: n=5). **Right**: Female mice (WT: n=3, Ud6-m-/p+: n=3, UG-m-/p+: n=2). Data points with Mean ± SEM of the group of pooled male and female mice (**left**), and male mice (**middle**) are plotted, *p<0.05, *** p<0.001, or as indicated by two-tail unpaired T test. Data points with Mean of the group of female mice (**right**) are plotted, and no significance is detected among genotypes in female mice.

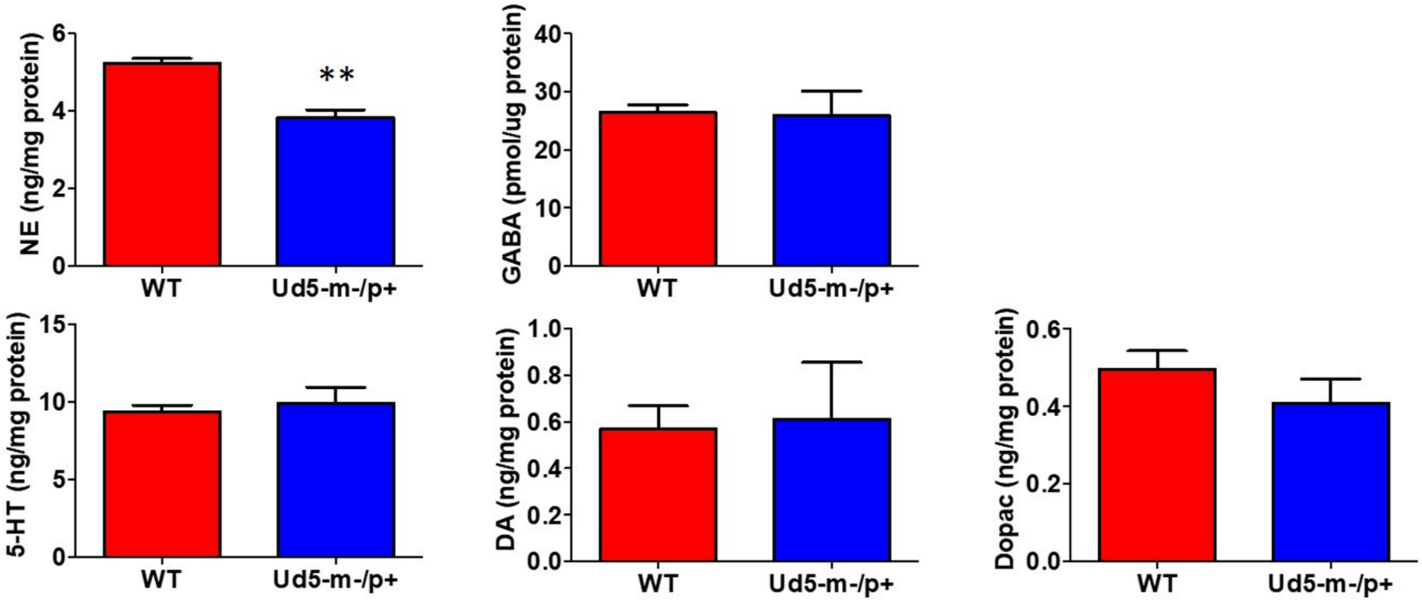


**Supplemental Figure 3. Levels of norepinephrine and other neurotransmitters in cerebellum of Ud5-m-/p+ mice.**

Mouse cerebellums were collected at CT 14 in DD (26 h in DD after release from LD 12:12). GABA and monoamine contents were analyzed by a LC/MS assay. Data are represented as the mean ± SEM (WT: N=5, Ud5-m-/p+: N=4). NE: norepinephrine, GABA: gamma-Aminobutyric acid, 5-HT: serotonin, DA: dopamine, Dopac: dihydroxyphenylacetic acid. ** $p < 0.01$ by two-tail unpaired T test.

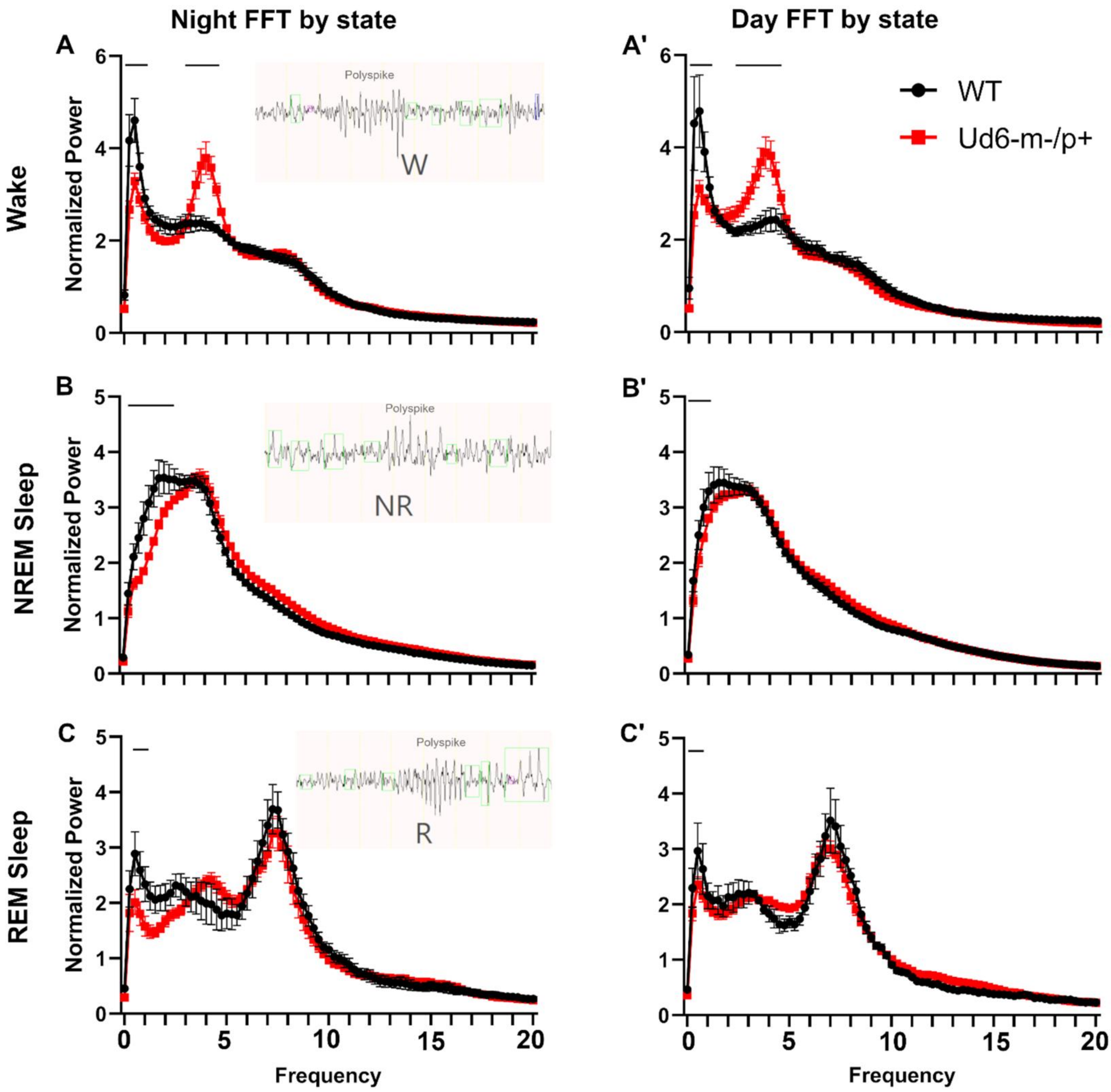


**Supplemental Figure 4. Spectral analysis of Wake, NREM sleep and REM sleep in LD condition.**

Ud6-m-/p+ (red) have lower delta (0.25-0.75Hz) power than WT in both the dark and light phase during (A,A') Wake, (B,B') NREM sleep and (C,C') REM sleep. Ud6-m-/p+ have more power in the range of high delta/low theta (3-4.25Hz). Polyspike episodes in this frequency were observed in Ud6-m-/p+ mice and insets of representative traces from each state are shown. Horizontal black lines indicate significance versus WT ($p<0.05$) following 2-way ANOVA and Bonferroni correction.

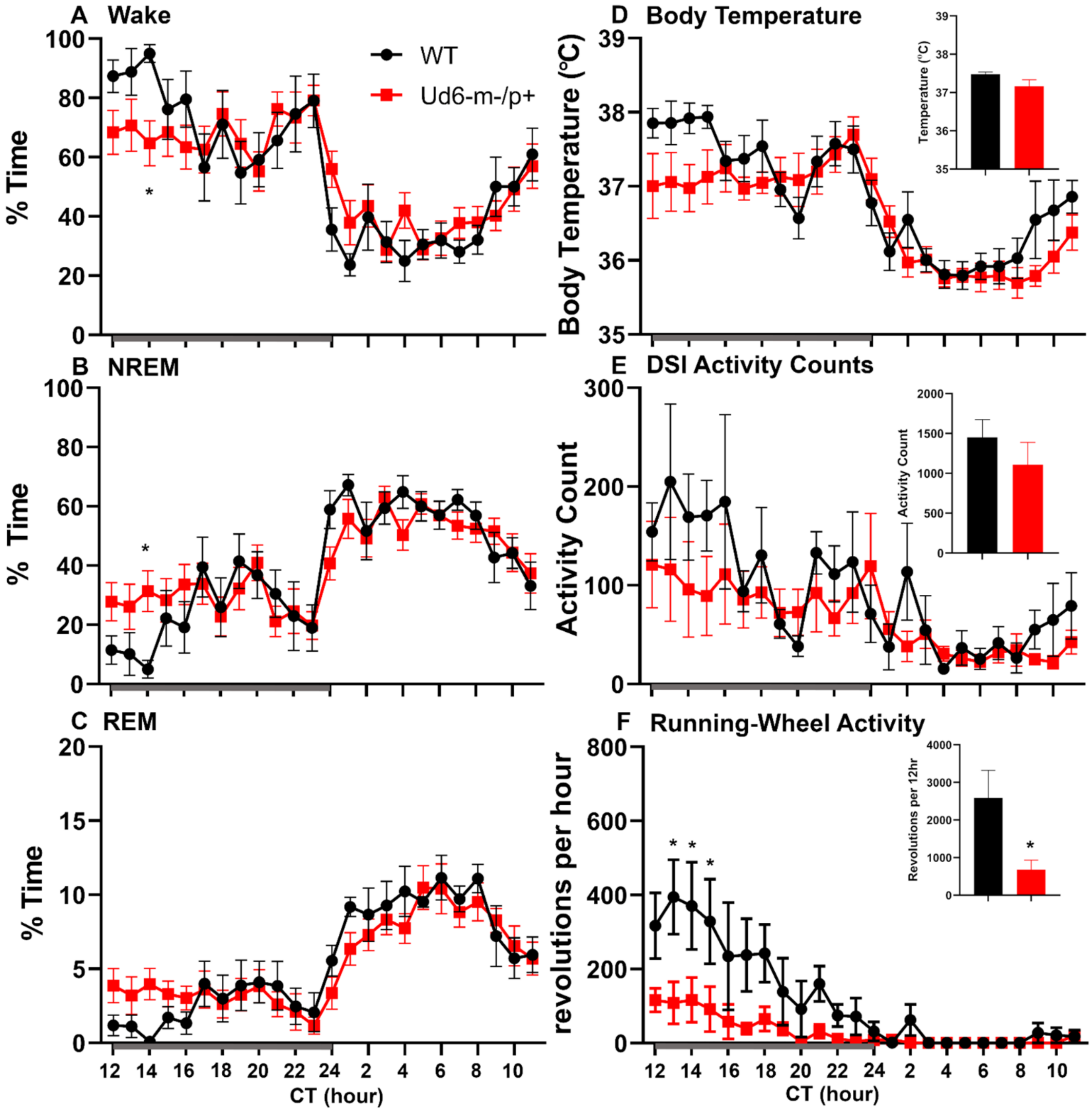


**Supplemental Figure 5. Constant dark conditions with access to a running wheel (DDRW).**

Under DDRW, amounts of wake (A), NREM sleep (B), and REM sleep (C) are similar in WT (black) versus Ud6-m-/p+ mice (red). During the initial active phase, Ud6-m-/p+ mice have lower body temperature (D), and reduced activity as recorded by DSI telemetry (E) and running-wheel revolutions (F). Active phase is indicated by the gray horizontal bar from CT12-24. Insets summarize the data for the 12-hour dark interval for WT (black) and Ud6-m-/p+ (red). Data are plotted as mean ± SEM. Asterisk (*) indicates a significant difference between Ud6-m-/p+ vs. WT mice at the $p<0.05$ level.

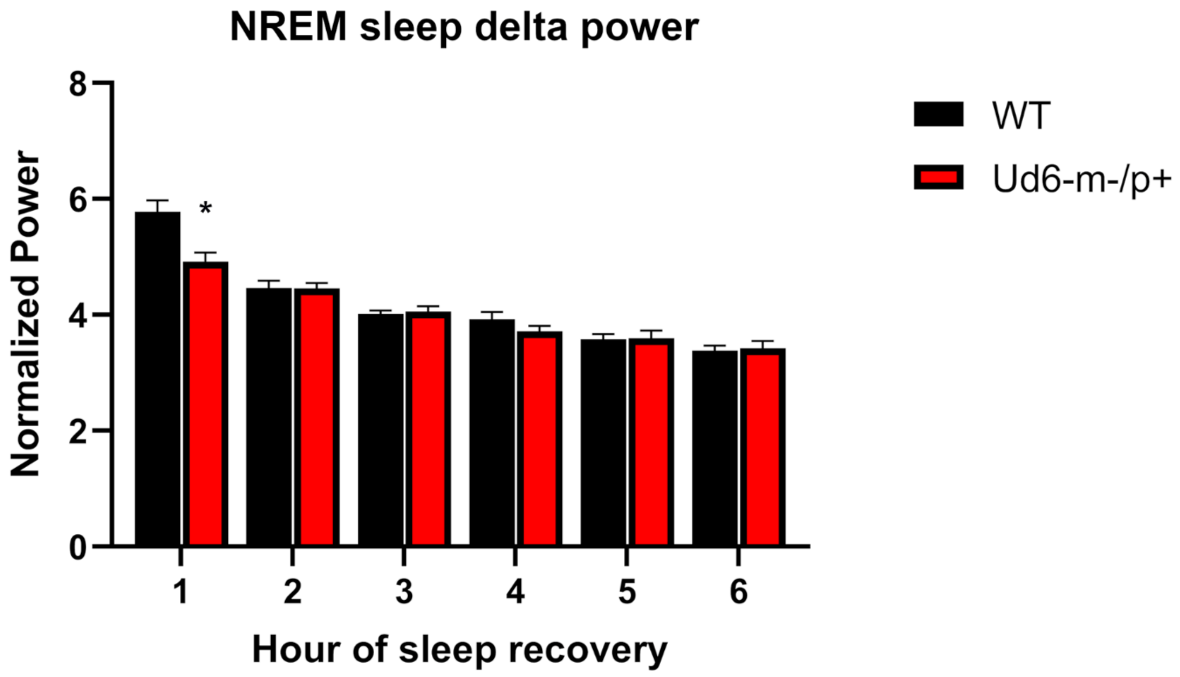


**Supplemental Figure 6. Spectral analysis of recovery sleep and wake following 6 hours of sleep deprivation.**

Ud6-m-/p+ (red) have lower NREM delta power than WT (black) in the first hour of sleep recovery following 6 hours of sleep deprivation. The area under the curve was calculated for each mouse with normalized power spectrum data in the range of 0.25 to 4 Hz (* vs. WT, $p<0.05$).

#### 2. Supplemental Data; Tables S1-S2

| *Supplemental Table 1* | | **amount (%)** | | ***# of bouts*** | | ***bout duration (sec)*** | |
|---|---|---|---|---|---|---|---|
| | | **Subjective night** | **Subjective day** | **Subjective night** | **Subjective day** | **Subjective night** | **Subjective day** |
| *Wake* | WT | 74 ± 1.8 | 36.1 ± 1.4 | 79 ± 8.9 | 152 ± 9.3 | 447.7 ± 61.3 | 105.3 ± 4.2 |
| | Ud6-m-/p+ | 68.4 ± 3.3 | 41 ± 2 | 106 ± 11.7 | 150 ± 7.2 | 347.6 ± 75.4 | 120 ± 8.2 |
| *NREM* | WT | 23.6 ± 1.7 | 55.2 ±1.2 | 74 ± 9.3 | 158 ± 8.5 | 146.5 ± 14.2 | 157.8 ± 11 |
| | Ud6-m-/p+ | 28.5 ± 3 | 51.3 ± 1.9 | 103 ± 12.4 | 157 ± 6.9 | 124.6 ± 6.8 | 143.9 ± 7.8 |
| *REM* | WT | 5.5 ± 1.3 | 11.2 ± 0.9 | 17 ± 1.82 | 55.3 ± 3.4 | 62.6 ± 4.2 | 67.7 ± 3.4 |
| | Ud6-m-/p+ | 3 ± 0.4 | **7.7 ± 0.4*** | 26 ± 3.5 | 59 ± 3 | 57.3 ± 4.5 | **56.6 ± 2.4*** |

Sleep data under DDRW conditions. * vs. WT, p<0.05 unpaired T-test.

**Supplemental Table 1. 12-hr summary of sleep under constant dark with access to a running wheel condition (DDRW).** Data are means ± SEM. Asterisk (*) indicates a significant difference between Ud6-m-/p+ vs. WT mice at the $p<0.05$ level.

| ***Supplemental Table 2*** | | **amount (sec)** | ***# of bouts*** | ***bout duration (sec)*** |
|---|---|---|---|---|
| | | **Recovery 6 hours** | **Recovery 6 hours** | **Recovery 6 hours** |
| *Wake* | WT | 6738.8 ± 481.6 | 74.1 ± 3.8 | 107.9 ± 5.2 |
| | Ud6-m-/p+ | 7174.2 ± 256.5 | 74.7 ± 3.4 | 124.2 ± 7.6 |
| *NREM* | WT | 12770 ± 403.7 | 75.8 ± 3.5 | 178.3 ± 11.3 |
| | Ud6-m-/p+ | 12275.8 ± 228.2 | 79.4 ± 2.8 | 161.5 ± 7.2 |
| *REM* | WT | 2091.3 ± 93.4 | 33.5 ± 2.3 | 72.5 ± 5 |
| | Ud6-m-/p+ | 2150 ± 108.9 | 35.7 ± 1.5 | 63 ± 2 |

Sleep data following 6hr sleep deprivation.

**Supplemental Table 2. 6-hr summary of sleep following sleep deprivation**. Ud6-m-/p+ vs. WT mice had similar responses to 6 hours of sleep deprivation. Data are means ± SEM.
